# Prolonged T cell – DC macro-clustering within lymph node microenvironments initiates Th2 cell differentiation in a site-specific manner

**DOI:** 10.1101/2023.07.07.547554

**Authors:** Miranda R. Lyons-Cohen, Elya A. Shamskhou, Michael Y. Gerner

## Abstract

Formation of T helper 2 (Th2) responses has been attributed to low-grade T cell stimulation, yet how large-scale polyclonal Th2 responses are generated *in vivo* remains unclear. Here, we used quantitative imaging to investigate early Th2 differentiation within lymph nodes (LNs) following cutaneous allergen administration. Contrary to current models, Th2 differentiation was associated with enhanced T cell activation and extensive integrin-dependent ‘macro-clustering’ at the T-B border, which also contrasted clustering behavior seen during Th1 differentiation. Unexpectedly, formation of Th2 macro-clusters within LNs was highly dependent on the site of skin sensitization. Differences between sites were driven by divergent activation states of migratory cDC2 from different dermal tissues, with enhanced costimulatory molecule expression by cDC2 in Th2-generating LNs promoting T cell macro-clustering and cytokine sensing. Thus, generation of dedicated priming micro-environments through enhanced costimulatory molecule signaling initiates the generation of Th2 responses *in vivo* and occurs in a skin site-specific manner.

## Introduction

Upon activation, naïve CD4 T cells differentiate into distinct helper cell lineages with specific effector functions tailored to eliminate different classes of pathogens. Th2 cells provide defense against parasitic helminth infections and promote tissue repair, but when inappropriately activated, can cause allergic disease or asthma^1^. Much work has gone into understanding the cellular and molecular mechanisms driving early Th2 differentiation, collectively resulting in the quantitative and qualitative models, which are also somewhat divergent from how other T helper cell lineages are thought to be generated^2, 3^.

The quantitative model posits that the signal strength sensed during initial T cell activation is a major factor regulating T helper cell polarization. Th2 differentiation has been suggested to involve decreased T cell receptor (TCR) signaling, either through reduced TCR affinity, lower levels of peptide MHC (pMHC) complexes presented by antigen presenting cells (APCs), or through limited sensing of costimulatory molecules^2–4^. Reduced signaling is thought to decrease the longevity of T cell – DC interactions, thus minimizing the ability of T cells to respond to inflammatory cytokines from DCs and ultimately promoting an endogenous program of Th2 polarization^2, 5, 6^. Confounding this model however is the notion that generally all *in vivo* responses involve polyclonal T cell populations with diverse TCR affinities, yet Th2 cells are not generated in all inflammatory contexts. Additionally, enhanced exposure to costimulatory molecules has been positively and not negatively associated with Th2 differentiation^7–12^. It is also not clear how low-grade stimulation elicits large scale *in vivo* Th2 responses as observed during helminth or allergen exposure, especially given that both result in maturation of cDCs and significant costimulatory molecule expression^13–16^.

In addition to quantitative signal strength -based factors, qualitative sensing of polarizing cytokines is important for T cell differentiation *in vivo.* However, unlike other helper cell lineages, Th2-promoting cytokines do not appear to operate in a typical ‘signal 3’ fashion through production by APCs^17^. Interleukin (IL) -4 is critical for Th2 differentiation both *in vitro* and *in vivo*, but this cytokine is not produced by cDCs and the exact cellular source/s of IL-4 in LNs remains ill-defined^1, 17^. Notably, recently activated T cells can produce IL-4 after TCR stimulation independently of the Th2 lineage-defining transcription factor, Gata3, and it has been suggested that paracrine delivery of IL-4 between activated T cells is sufficient for Th2 differentiation^18–21^. Similarly, T cell derived IL-2, also produced downstream of T cell activation, is necessary for Th2 response formation *in vivo*^21, 22^, and this cytokine is again delivered via paracrine exchange between activated T cells and not provided by APCs^23, 24^.

Moreover, not all cDC populations have equivalent capacities to induce Th2 responses. Following barrier tissue damage, locally released alarmins induce the activation of cDC2s, including cells expressing CD301b and variegated levels of Sirpα/CD11b expression^13, 14, 25^. Activated cDC2s in turn migrate into draining LNs, where they can induce Th2 responses^13, 14, 25–30^. cDC2s, however, are also highly plastic and based on the nature of the stimulus can generate diverse helper lineages, including T follicular helper (Tfh), Th1, and Th17 cells^31–34^, and the exact molecular mechanisms of how these cells promote Th2 responses during type-II inflammation remain unknown. cDC1s on the other hand constitutively secrete IL-12 and inhibit Th2 responses, instead promoting Th1 and CD8 T cell immunity^35, 36^. Optimal Th2 differentiation thus likely involves both selective engagement with appropriately activated cDC2 and avoidance of cDC1 populations. How such selectivity is achieved *in vivo* is unknown, although recent quantitative imaging studies demonstrated that different cDC subsets are non-equivalently spatially distributed within LNs, which could allow for preferential engagement vs. avoidance of specific DC subsets by responding T cells in distinct tissue compartments^37^. Indeed, during type-I inflammation, the spatial positioning of specific innate subsets, including monocytes and activated cDCs, establishes the formation of dedicated microenvironments in the deep T cell zone to generate effector Th1 and CD8 T cell responses^38, 39^. In contrast, CD301b^+^ cDC2s predominantly localize at the T/B border, a location where early Th2 cells have also been previously noted^25, 40–42^. This suggests an additional underexplored spatial component of Th2 differentiation in which LN microenvironments populated by appropriately instructed myeloid subsets drive T cell differentiation towards distinct helper lineages.

Here, we used quantitative microscopy to investigate the early stages of *in vivo* Th2 differentiation after cutaneous administration of the allergen, papain, and other type-II stimuli, as well as compared these responses to Th1 differentiation with TLR agonist immunization. In contrast to the predicted limited cellular activation in Th2 settings, we observed enhanced T cell signaling and extensive clustering, here termed ‘macro-clustering’, of early differentiating Th2 cells which primarily occurred near the T-B border of the LN paracortex. Macro-clustering was integrin-mediated and was associated with enhanced local cytokine signaling, suggesting that the spatial proximity of activated T cells enabled optimized cytokine exchange for Th2 differentiation. Enhanced T cell signaling and macro-clustering were also distinct from that seen with adjuvant induced Th1 responses. Surprisingly, formation of Th2 responses was highly dependent on the specific skin site of type-II agonist administration, with footpad-delivered stimuli eliciting markedly reduced Th2 responses as compared to other skin sites, but without compromised ability to elicit Th1 differentiation across different tissues. Th2 differentiation in LNs was driven by migratory cDC2s, and the observed site specific generation of Th2 responses enabled a direct comparison of key cDC2 properties necessary for Th2 differentiation using the same agonist. This revealed divergent activation states for migratory cDC2 emigrating from different skin sites and a critical role for enhanced costimulatory molecule expression coupled with low levels of pMHC-II to drive T cell macro-clustering, cytokine signaling, and Th2 differentiation. Collectively, our findings argue that enhanced costimulation and integrin driven signaling coupled with low grade TCR stimulation promotes T cell macro-clustering and generates LN microenvironments which drive Th2 response formation *in vivo*. Our data also support the emerging notion that generation of T cell responses is heavily impacted by upstream barrier tissues^43, 44^, raising fundamental questions on the mechanisms leading to divergent responses among skin sites and having clear implications for allergic disease development.

## Results

### Generation of Th2 microenvironments in skin draining LNs

To examine the early processes governing *in situ* Th2 differentiation, we crossed Ovalbumin (OVA)-specific TCR-transgenic OT-II mice to IL-4 mRNA reporters^45^ to generate 4get-GFP OT-II mice (4get-GFP.OT-II) on a CD45.2 congenic background. We then adoptively transferred naïve 4get-GFP.OT-II CD4 T cells into CD45.1^+^ recipient B6 mice, administered papain plus OVA intradermally in the ear pinnae one day later, and examined the localization and phenotype of activated OT-II cells in auricular (Au) draining LNs 2-3 days after immunization using quantitative multiparameter microscopy. Papain is a cysteine protease and a clinically relevant allergen in humans and mice that drives robust induction of type-II immunity after cutaneous administration^25^. CD62L blocking antibody was also administered 6 hours after immunization to minimize the impact of asynchronous activation of naïve T cells recruited into LNs late into the response^46^. We observed formation of extensive macro-clusters of OT-II cells located primarily at the T-B border of draining LNs (Figure 1A), and the clustered cells expressed high levels of IRF4 and Ki67, indicative of T cell activation and proliferation (Figure 1A, 1B region 1 inset). Cells within the clusters also expressed high quantities of the Th2 lineage-defining transcription factor, Gata3, and were positive for the 4get-GFP reporter signal, indicating IL-4 mRNA transcription. In contrast, fewer activated OT-II cells outside the macro-clusters expressed Gata3 and 4get-GFP (Figure 1A, 1B region 2 inset), suggesting that the macro-clusters represented regions where T cells underwent their earliest stages of Th2 lineage commitment detectable with this approach. Large Th2 macro-clusters at the T-B border were also observed when examining endogenous CD4 T cell responses after papain inoculation of 4get-GFP mice, indicating that this macro-clustering phenomenon was occurring during polyclonal endogenous T cell activation, and was not an artifact of high precursor frequency after adoptive transfer (Figure S1A). Moreover, macro-clusters of IRF4, Gata3 and 4get-GFP-expressing polyclonal Th2 cells were also found in mesenteric LNs 6 days post infection with the helminth *Nippostrongylus brasiliensis* (N.b.), a timepoint when the parasite establishes robust infection in the intestine and induces Th2 priming^47^ (Figure S1B).

**Figure 1.**
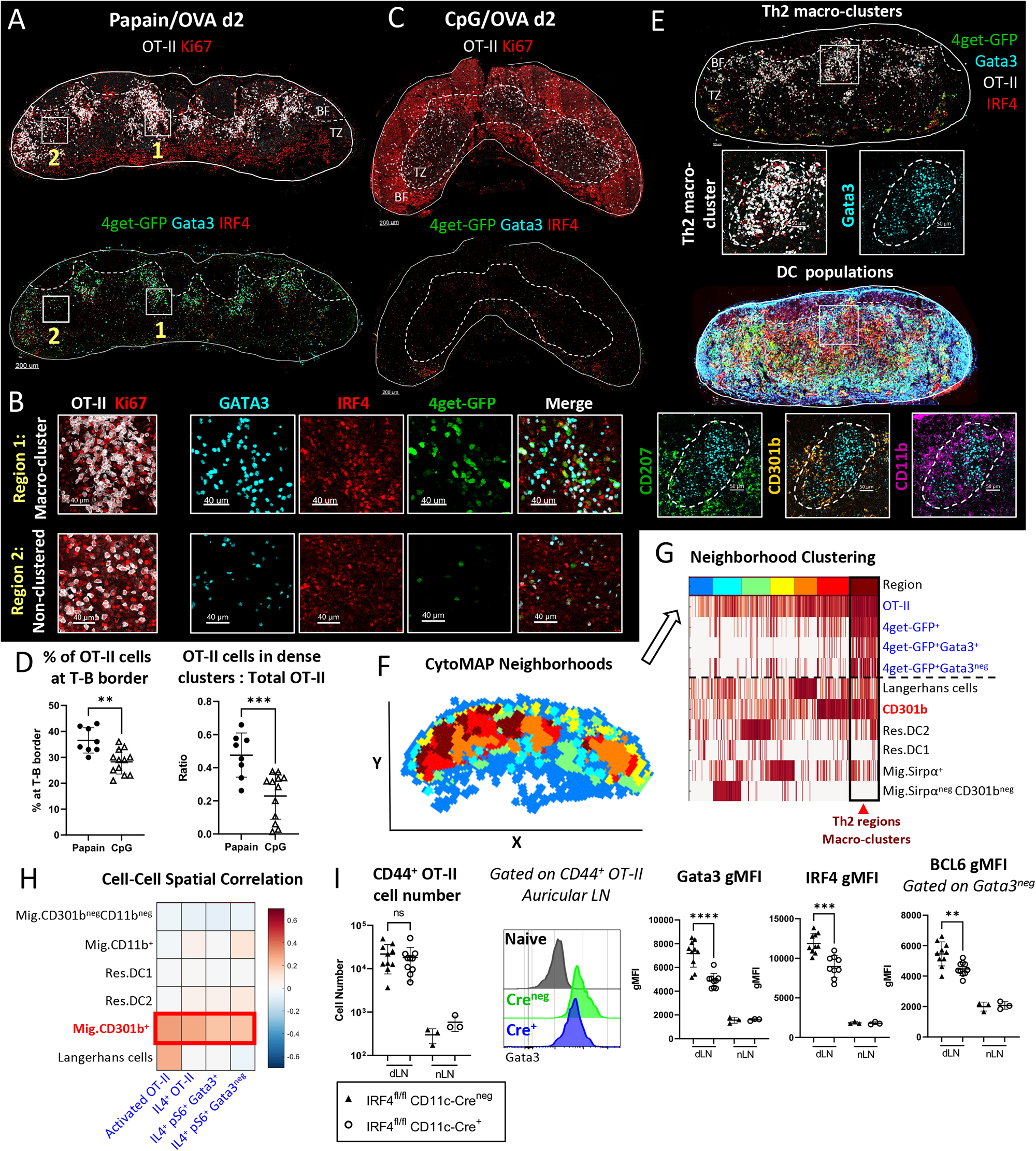
Generation of Th2 microenvironments in draining LNs. A-H) Naïve Ly5.2 4get.OT-II T cells were transferred to Ly5.1 mice and injected with the indicated adjuvants plus OVA in the ear pinnae. Mice were treated intraperitoneally with α-CD62L blocking antibody 6 hours post immunization and dLNs were harvested and assessed by histo-cytometry at 48 hours. A-B) Representative images depicting OT-II macro-clustering at the T-B border in Papain OVA immunized dLNs with B) macro-clustered and non-clustered regions highlighted in insets with the indicated markers shown. C) Representative images depicting OT-II responses in dLNs after CpG OVA immunization. D) Histo-cytometry analyses of OT-II localization with respect to the T-B border and the ratio of macro-clustered vs. total OT-II are shown. E-H) dLN tissue sections were analyzed for association of Th2 cells with myeloid cells. E) Representative image demonstrating Th2 macro-clusters and different myeloid cell markers. F-H) Spatial distribution analysis of myeloid cells and activated (IRF4^+^Ki67^+^) T cell subsets was performed using CytoMAP. F) 50μm raster scan neighborhoods of a representative dLN were plotted on a X,Y positional plot and color-coded using neighborhood clustering, as shown in panel G. G) Heatmap demonstrating the cellular composition for the different neighborhood clusters (color code depicted at the top). Th2 regions (dark red) enriched with CD301b^+^ DCs are highlighted (arrowhead). H) Cell-cell spatial correlation of the indicated T cell subsets and myeloid cell populations. I) Naïve Ly5.1 OT-II T cells were transferred into CD11c-Cre^+^ IRF4^fl/fl^ or CD11c-Cre^neg^ IRF4^fl/fl^ Ly5.2 mice and immunized with papain and OVA in the ear pinnae, treated with α-CD62L blocking antibody at 6 hours, and dLNs were harvested and assessed by flow cytometry at 48 hours. I) CD44^+^ OT-II cell number is shown with representative histograms of Gata3 gMFI of CD44^+^ OT-II cells, IRF4 gMFI, and BCL6 gMFI on Gata3^neg^ CD44^+^ OT-II cells. BF = B cell follicles; TZ = T cell zone. Dashed lines represent T-B border. Figures A-D are representative of 5 independent experiments. Figures E-I are representative of 3 independent experiments.

As a comparison, we examined responses after OVA plus CpG immunization, which promotes Th1 differentiation^38^. CpG OVA immunization elicited robust OT-II activation and proliferation indicated by Ki67 expression, but these T cells had undetectable Gata3 and 4get-GFP expression, corresponding to lack of Th2 differentiation in these settings (Figure 1C). In contrast to papain induced responses, CpG immunization did not elicit the formation of dense T cell macro-clusters and instead generated much smaller cell clusters which were more diffusely distributed throughout the T cell zone and the outer LN paracortex, consistent with past findings on behavior of CD4 T cells during Th1 differentiation^38, 39^ (Figure 1C-D).

Previous studies visualizing IL-4 producing cells in LNs at late time points have been conflated by detection of IL-4 producing Tfh cells within B cell follicles^48, 49^, so we examined Tfh markers on the responding 4get-GFP^+^ T cells 2-3 days after papain OVA treatment. We found that most of the 4get-GFP^+^ cells displayed high levels of the high affinity IL-2 receptor, CD25, and low levels of CXCR5 and PD-1 staining (Figure S1C), indicating early effector T cell and not Tfh differentiation^50–52^. Similarly, endogenous activated (CD44^+^Ki67^+^) Gata3^+^ CD4 T cells expressed CD25 and lacked BCL6 expression, while a separate population of CD25^neg^ cells co-expressed Gata3 and BCL6, indicating bifurcation of effector lineages (Figure S1D)^53–56^. To verify that IL-4 mRNA competent 4get-GFP^+^ T cells also produced IL-4 protein, we examined responses in KN2^+/-^ reporter mice which express human CD2 (huCD2) on the surface of T cells actively producing IL-4 protein^57^. We found abundant huCD2 expression on endogenous responding Gata3^+^ CD44^+^Ki67^+^ CD4 T cells suggesting active IL-4 protein production (Figure S1E).

To investigate which specific myeloid cell population/s were associated with the Th2 macro-clusters, we next co-stained sections of papain-immunized auricular draining LNs with various innate cell markers and used histo-cytometry and CytoMAP to analyze myeloid cell composition and distribution^42, 58^. Neighborhood clustering analysis identified distinct LN regions populated by different myeloid cell subsets (Figure 1E-G). As previously reported, migratory cDC2s, including CD11b^+^CD301b^+^ and CD11b^+^CD301b^neg^ subsets were predominantly localized in the outer T zone regions, with CD301b^+^ cDC2s localized at the T-B border and in close proximity to the Th2 macro-clusters^25, 40, 59^ (Figure 1E). In contrast, CD11b^+^CD207^+^ Langerhans cells were predominantly distributed within the deeper T cell zone and appeared spatially segregated from the Th2-dense regions (Figure 1E-G). CytoMAP spatial correlation analysis across multiple LNs confirmed these observations, demonstrating that CD301b^+^ cDC2s were the dominant myeloid cell subset spatially correlated with Th2 cells, while neighborhood clustering identified distinct regions with preferential CD301b^+^ cDC2s and Th2 enrichment (Figure 1F-H). Together these data demonstrate the formation of Th2 microenvironments defined by macro-clustering of early differentiating Th2 cells and migratory cDC2 subsets.

cDC2s have been previously demonstrated to drive Th2 polarization, but not T cell proliferation, after papain immunization^25, 60^. To examine the requirement of migratory cDC2s for Th2 responses in our model, we examined OT-II cell responses in IRF4^fl/fl^ CD11c-Cre^+^ mice which exhibit impaired cDC2 migration from peripheral tissues into LNs, while keeping other cDC populations intact^27, 30^ (Figure S1F). Indeed, as compared to Cre^neg^ littermate controls, loss of migratory cDC2 in IRF4^fl/fl^ CD11c-Cre^+^ animals significantly reduced Gata3 and IRF4 expression in the responding OT-II cells without altering their clonal expansion, suggesting abrogated Th2 differentiation but not priming (Figure 1I). cDC2s have also been reported to promote Tfh differentiation^31, 32, 61^, and we noted reduced BCL6 expression in responding CD44^+^ Gata3^neg^ OT-II cells in IRF4^fl/fl^ CD11c-Cre^+^ mice (Figure 1I), suggesting that migratory cDC2s mediate the initiation of both Th2 and Tfh responses. Altogether, these data indicate that cutaneous papain administration into the ear pinnae induces the formation of Th2-promoting microenvironments at the T-B border composed of macro-clusters of highly activated, early differentiating Th2 cells and CD301b^+^ migratory cDC2s, which drive Th2 response induction, and that this clustering behavior is also distinct from that observed during Th1 differentiation after TLR agonist immunization.

### Th2 macro-clustering and differentiation is site-specific

Cutaneous exposure to allergens and associated antigens can occur in distinct anatomical locations. Surprisingly, when administering papain OVA into distinct skin sites, we observed major differences in Th2 response induction within the corresponding skin draining LNs. As above, auricular LNs draining ear pinnae generated extensive early Th2 macro-clustering at the T-B border (Figure 2A). In contrast, the equivalent dose of papain and antigen administered in the footpad led to minimal Th2 differentiation in the draining brachial (Br) LNs (Figure 2B-C). Instead of macro-clustering at the T-B border, most OT-II T cells in brachial LNs were more homogenously distributed and localized in smaller clusters throughout the T cell zone, and expressed significantly less Gata3 and 4get-GFP as detected by histo- and flow cytometry (Figure 2A-E). Decreased Th2 response formation was not due to general lack of T cell activation, as following footpad immunization brachial LN OT-II T cells expressed abundant Ki67 and underwent equivalent or even greater levels of early proliferation as compared to those in auricular draining LNs (Figure 2B, E).

**Figure 2.**
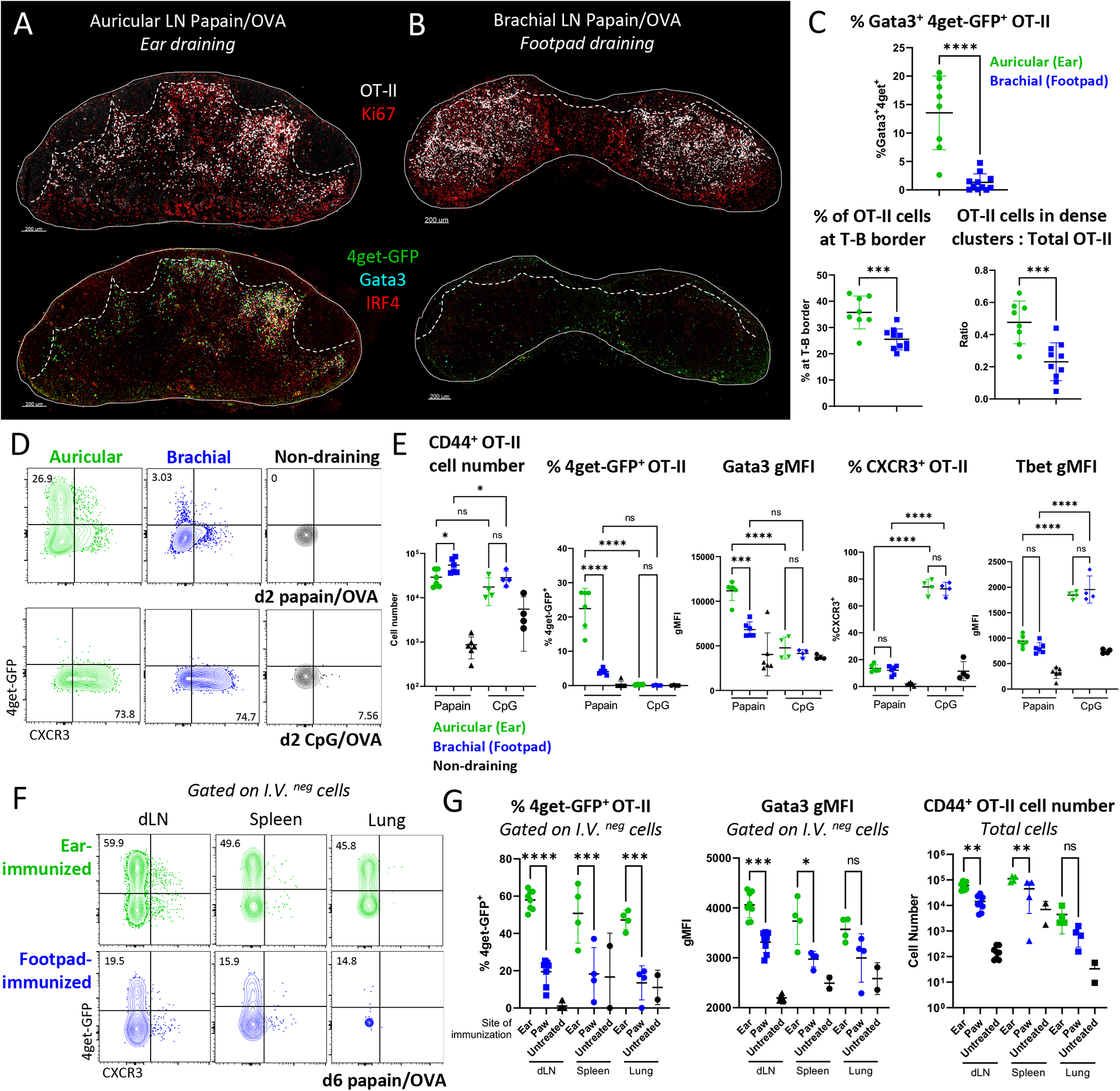
Th2 macro-clustering and differentiation is site-specific. A-E) Naïve Ly5.2 4get.OT-II cells were transferred to Ly5.1 mice and injected with the indicated adjuvant plus OVA in the ear pinnae or front footpad, treated intraperitoneally with α-CD62L 6 hours post immunization, and dLNs were harvested and assessed by histo-cytometry and flow cytometry at 48 hours. Representative images depicting OT-II macro-clustering and expression of the indicated markers in A) auricular vs B) brachial dLNs. C) Histo-cytometry analysis of the percent activated OT-II co-expressing 4get-GFP and Gata3, OT-II localization, and the ratio of macro-clustered vs. total OT-II in the indicated LNs. D-E) Representative plots and quantification of CD44^+^ OT-II cell number, frequency of 4get-GFP^+^ and CXCR3^+^ cells, and Gata3 and Tbet gMFI of CD44^+^ OT-II cells. F-G) 0.5×10^6^ naïve Ly5.2 4get.OT-II cells were transferred to Ly5.1 mice then injected with papain OVA in the ear pinnae or front footpad. 6 days later, cells were labeled intravenously, and dLNs, spleen, and lung were harvested and assessed by flow cytometry. Representative plots and quantification of frequency of 4get-GFP^+^ and Gata3 gMFI of I.V.^neg^ CD44^+^ OT-II cells and total CD44^+^ OT-II cell number are shown. Au = auricular; Br = brachial; IV = intravenous. Dashed lines represent T-B border. A-E are representative of 5 independent experiments. F-G are representative of 2 independent experiments.

Site-specific differences between the ear and footpad were maintained for at least 6 days and across peripheral organs, with significantly reduced frequencies of effector Th2 OT-II cells also found in the parenchyma of the lung and spleen, suggesting that divergent T cell responses were maintained even after OT-II cells migrated out of the original site of priming (Figure 2F-G). Despite the similar initial clonal bursts, by day 6, OT-II cells primed in auricular draining LNs exhibited increased expansion as compared to those primed in brachial draining LNs and greater numbers of OT-II cells disseminated to the spleen and lungs (Figure 2G). At these later time points, the auricular draining LNs also exhibited increased frequency of CXCR5^+^PD-1^+^ Tfh OT-II cells, and the cells also expressed a greater amount BCL6, suggesting that both Th2 and Tfh responses were linked to the specific skin site of immunization (Figure S2A). Site-specific Th2 differences were also observed for endogenous CD4 T cells in KN2^+/-^ reporter mice, with a significant reduction of IL-4 producing Gata3^+^ CD4 T cells in footpad draining brachial LNs (Figure S2B). Th2 response differences across immunization sites were also observed in Balb/c mice, indicating that this phenomenon was conserved across mouse strains with differential abilities to drive type-II immunity^62^ (Figure S2C), as well as after OVA Alum immunization in endogenous T cell and OT-II settings (Figure S2D-E). We next examined Th1 differentiation settings after CpG OVA immunization in distinct cutaneous tissues. In stark contrast to type-II inflammation settings, CpG OVA resulted in equivalent expansion, Tbet expression, and CXCR3 expression in responding OT-II cells in both auricular and brachial LNs, indicating comparable Th1 response formation between the different sites (Figure 2E). Equivalent Th1 responses across these distinct LNs were also previously observed with other type-I adjuvants^38^.

Non-equivalent Th2 responses in different skin draining LNs may result from intrinsic differences between the LNs, differential lymphatic drainage of antigens and agonists, or fundamental differences in how distinct cutaneous sites program local immune responses. To investigate whether brachial LNs have an inherent defect in mounting Th2 immunity, we immunized mice with papain OVA in the dorsal skin of the flank, which targets a different skin dermatome but antigen and agonist also drain into brachial LNs. Although generating more heterogeneous responses as compared to footpad inoculation, likely due to more diffuse agonist dispersal across the subcutaneous tissue compartment, flank injection resulted in relatively normal Th2 induction in brachial LNs, which was more analogous to that found in auricular LNs with ear immunization (Figure S2F-G). We also tested other cutaneous sites, including the hind paw and the tail base which drain the popliteal and inguinal LNs, respectively. Similar to brachial LN responses with forepaw injection, hind paw inoculation also elicited limited Th2 responses in the draining popliteal LNs, with decreased Gata3 expression by activated OT-II cells. In contrast, tail base administration of papain OVA generated heightened Th2 responses within the inguinal draining LNs (Figure S2F-G). We next examined whether site-specific Th2 response differences would be maintained in a model which does not involve abundant lymphatic drainage induced by injection. For this, we painted the ear vs. footpad skin with dibutyl phthalate (DBP) to induce a model of Th2-driven contact hypersensitivity in which antigen and adjuvant are administered epicutaneously and not intradermally^63^. Although DBP painting did not induce as potent of proliferative responses as papain injection, major differences in Th2 differentiation were still observed across the draining LNs, with markedly increased Gata3 and IRF4 expression in responding CD4 T cells within auricular but not brachial draining LNs (Figure S2H).

Together, these findings indicate that distinct cutaneous sites have non-equivalent abilities to drive Th2 macro-clustering and Th2 differentiation, but that these site-specific differences do not necessarily extend to other helper cell lineages, such as Th1 responses. Moreover, non-equivalent responses between sites do not simply result from intrinsic differences between the lymphoid organs or are restricted to injection-based models, indicating a more generalizable divergence in the ability to drive Th2 responses across distinct skin sites.

### Costimulatory molecule expression is enhanced on migratory cDC2s derived from auricular LNs

Given the critical role of cDC2s in driving Th2 responses, we next examined the hypothesis that migratory cDC2 responses were non-equivalent for the distinct dermal tissues. To test this, we immunized mice with papain plus the fluorescent protein EαGFP into the forepaw or ear pinnae^64^, and examined antigen-bearing migratory DCs in the corresponding draining LNs. Albeit some variation was observed among individual samples, across multiple experiments we found no major differences in the number of total cDC2s, antigen-bearing EαGFP^+^ cDC2s, or in the amount of EαGFP captured by these cells between the sites (Figure 3A-C, S3A). The dominant antigen-bearing population in both draining LNs were the CD301b^+^ cDCs (Figure S3B), and their total number did not differ between draining LNs (Figure 3C). The overall composition of antigen-bearing cells was also largely similar across the LNs, albeit a modest increase in the frequency of antigen-bearing CD301b^+^ DCs and CD64^+^ cells and a corresponding decrease in CD301b^neg^CD11b^neg^ DCs in brachial LNs was noted (Figure S3B).

**Figure 3.**
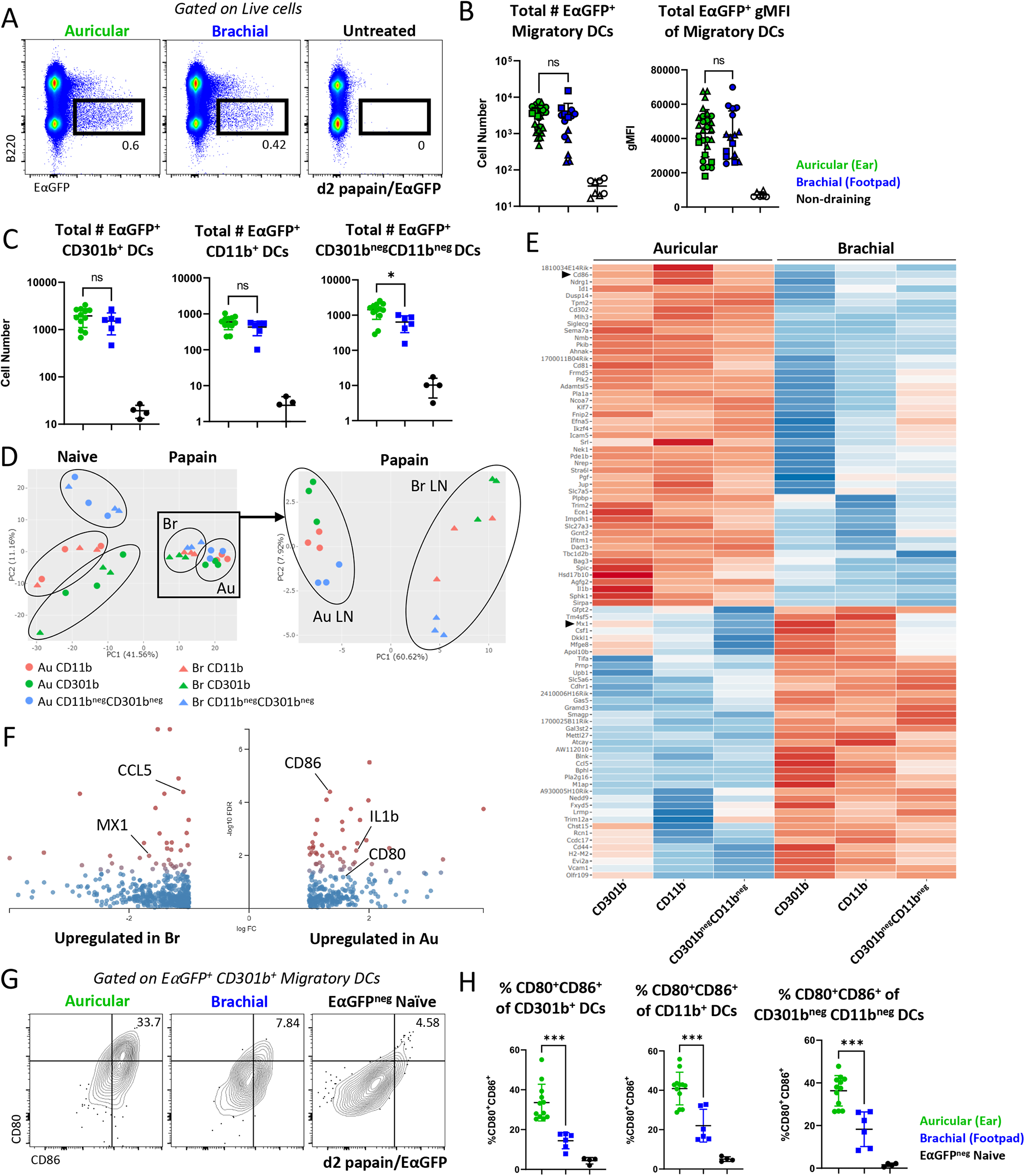
Costimulation is enhanced on DCs from auricular LNs. A-H) B6 mice were immunized in the ear pinnae and front footpad with papain plus EαGFP and the corresponding dLNs were harvested 48 hours later. A) Representative plots showing total EαGFP expression within live cells. B) Quantification of the number of EαGFP^+^ migratory DCs and EαGFP^+^ gMFI of migratory DCs. Symbol shapes represent independent experiments (three total). C) Quantification of total EαGFP^+^ cells for the indicated DC subsets. D-F) EαGFP^+^ DC populations were sorted from dLNs on day 2 for bulk RNA sequencing. DCs were sorted on Live, CD64^neg^, Lineage^neg^ (CD3, CD19, NK1.1), EαGFP^+^, MHC-II^Hi^ CD11c^Int^, XCR1^neg^, EpCAM^neg^, then sorted into CD11b^+^ CD301b^+^, CD11b^+^ CD301b^neg^, and CD11b^neg^ CD301b^neg^ populations. The same gates were used for EαGFP^neg^ DCs in naïve LNs. D) PCA plot of all samples (left) and papain immunized DC populations (right). E) Heatmap of the top 100 DEGs between antigen-bearing migratory DCs from auricular vs. brachial dLNs (FDR<0.05, 2x fold change expression). F) Volcano plot of DEGs between antigen-bearing migratory DCs from auricular and brachial dLNs. Red indicates a FDR<0.05. Genes with a log fold change greater than 2 are shown. G-H) EαGFP^+^ migratory DCs were assessed for surface CD80 and CD86 expression by flow cytometry. G) Representative flow plots for CD301b^+^ cDC2s and H) quantification for three indicated DC subsets are shown. Au = auricular; Br = brachial. A-C, and G-H are representative of 3 independent experiments. D-F are representative of 1 independent RNA sequencing experiment with n=3 per group.

We also examined the amount of pMHC-II complex presented on the cDC surface using the Y-Ae antibody, which recognizes the Eα peptide presented on I-Ab^64^. Surprisingly, we found that while papain induced robust antigen uptake by migratory cDCs, this was not associated with detectable pMHC-II complex on the cell surface (Figure S3C-D). Minor differences in the total number, but not frequency, of EαGFP^+^ Y-Ae^+^ migratory DCs were noted between auricular and brachial LNs after papain immunization (Figure S3D). In contrast to papain settings, EαGFP plus CpG immunization elicited both antigen uptake and robust surface pMHC-II expression by migratory cDCs (Figure S3C-D), demonstrating major differences in how type-I and type-II stimuli impact MHC-II antigen processing and presentation and suggesting that during *in vivo* type-II responses to papain, surface pMHC-II expression is relatively limited on migratory DCs. Of note, Eα peptide sequence analysis did not identify papain cleavage sites, indicating that the divergence in pMHC-II complex between papain and CpG conditions was not simply a result of protease mediated degradation of the Eα peptide.

To further interrogate potential differences among migratory cDCs from different cutaneous sources, we next sorted antigen-bearing (EαGFP^+^) migratory cDC2 subsets (MHC2^Hi^CD11c^Int^ CD301b^+/neg^CD11b^+/neg^) from auricular and brachial dLNs two days post papain plus EαGFP immunization, or from naïve mice, and performed bulk RNA sequencing. As expected, principal component analysis (PCA) of all samples demonstrated that the primary segregation was driven by the immunization state, with samples clustering based on whether they were obtained from naïve or papain immunized LNs, revealing large-scale transcriptional changes across all cDC populations from both tissues after papain immunization (Figure 3D left, S3E-F). Of note, IL-12b, which is constitutively expressed by migratory cDCs at steady state^35, 36^, was downregulated upon papain immunization (Figure S3F). When considering only the papain immunized samples, further PCA separation demonstrated that sample divergence on the PCA1 axis was dominantly driven based on the specific LN of origin, indicating major transcriptional differences between the antigen-bearing auricular-derived and brachial-derived cDCs, while also maintaining a secondary level grouping along the PCA2 axis based on the cell subset (Figure 3D right, S3E). Of the top 100 differentially expressed genes (DEGs) between sites, DCs from auricular LNs preferentially upregulated genes associated with activation and costimulation including CD80, CD86, PDL1, and PDL2, indicating increased cDC2 maturation (Figure 3E-F). Divergent expression of these molecules at the protein level was confirmed by flow cytometry, demonstrating that antigen-bearing migratory cDC2s in auricular LNs following ear pinnae immunization expressed significantly higher levels of CD80 and CD86 costimulatory molecules as compared to their antigen-bearing counterparts from brachial LNs following footpad immunization (Figure 3G-H). We also noted increased expression of PD-L1 and PD-L2 on auricular-derived cDC2s at the protein level (Figure S3G). In several subsets, cDC2 gene signatures in brachial LNs were also enriched in type 1 interferon (IFN) and viral sensing pathways such as expression of the interferon-stimulated gene, MX1, suggesting that additional differences exist between the sites (Figure 3F, S3H). In contrast, auricular gene signatures were enriched in cytokine-mediated and leukocyte proliferation signaling pathways. Together, these data suggest that while being relatively similar prior to inflammation, cDC2s migrating into LNs from different cutaneous sites exhibit marked differences at the transcriptional and protein levels, and in particular for costimulatory molecule expression.

### Site-specific T cell response differences are mediated through non-equivalent expression of costimulatory molecules by migratory cDCs

Costimulation synergizes with TCR signaling to drive optimized T cell activation and has been implicated in Th2 differentiation^4, 7–11, 65, 66^. Given the costimulatory molecule expression differences on DCs between the sites, we investigated whether activation of signaling pathways downstream of TCR and costimulation were also nonequivalent in activated T cells in different draining LNs. Indeed, we found significantly higher expression of the AP-1 transcription factors, BATF and IRF4, in activated OT-II T cells within the ear draining auricular LNs, indicating non-equivalent engagement of the TCR / costimulatory molecule signaling platform (Figure 4A-B, 2A-B). Expression of the transcription factors BATF and IRF4 in T cells has been previously associated with Th2 differentiation and IL-4 production^67–75^, and we observed a strong positive correlation of these molecules with each other and Gata3 and 4get-GFP expression (Figure 4C). Image-based analysis also demonstrated increased phosphorylation of the mTOR signaling protein S6 (pS6), also downstream of TCR / costimulation platform, in T cells within auricular draining LNs (Figure 4D). Unexpectedly, in contrast to type-II inflammation, CpG OVA immunization promoted overall reduced levels of pS6 and IRF4 as compared to papain within auricular LNs, and equivalent site-specific expression across both LN sites (Figure 4A, D). This indicates that Th2 differentiation is associated with relatively enhanced and not reduced T cell activation, and that site specificity in T cell response induction does not extend to all T helper cell lineages.

**Figure 4.**
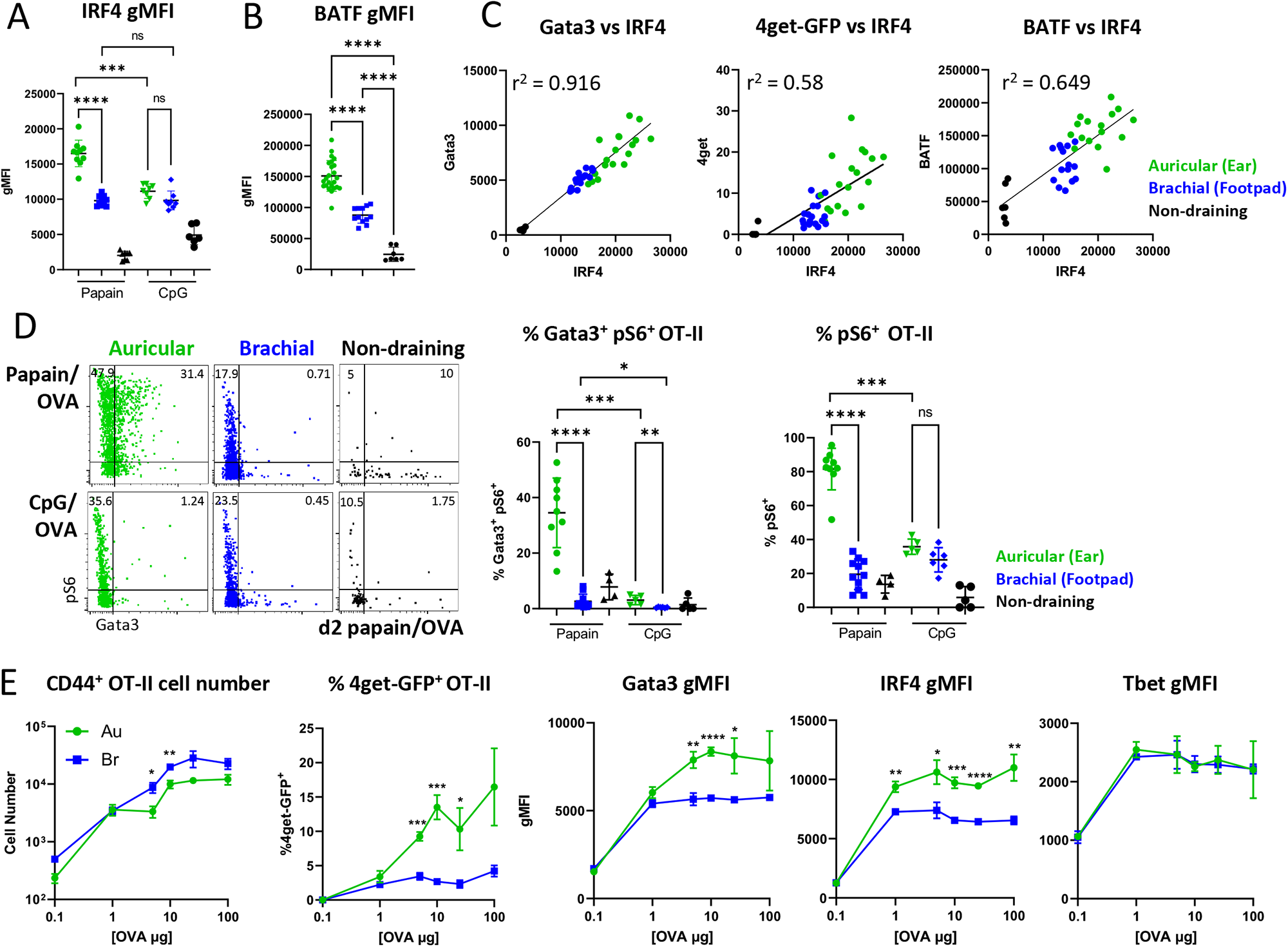
Non-equivalent expression of costimulatory molecules by migratory cDC2s drives divergent T cell activation states. A-D) Naïve Ly5.2 4get.OT-II cells were transferred to Ly5.1 mice, injected with papain OVA or CpG OVA in the ear pinnae or front footpad, treated intraperitoneally with α-CD62L 6 hours post immunization, and dLNs were harvested and assessed by histo-cytometry or flow cytometry at 48 hours. A-C) Expression of IRF4 and BATF on activated CD44^+^ OT-II cells and correlation plots with Gata3 gMFI and frequency of 4get-GFP^+^ cells across different dLNs is shown. D) Representative histo-cytometry plots and quantification of pS6 and Gata3 expression in activated Ki67^+^IRF4^+^ OT-II T cells. E) Naïve Ly5.2 4get.OT-II cells were transferred to Ly5.1 mice and injected with a fixed concentration of papain and increasing concentrations of OVA in the ear pinnae or front footpad. Mice were treated intraperitoneally with α-CD62L 6 hours post immunization and dLNs were harvested and assessed by flow cytometry at 48 hours. CD44^+^ OT-II cell number, frequency of 4get-GFP^+^ cells, and Gata3, IRF4, and Tbet gMFI expression on CD44^+^ OT-II cells are shown. Data are representative of at least 3 independent experiments.

Together, these findings suggested that after papain immunization, monoclonal CD4 T cells within ear draining auricular LNs experienced increased overall stimulation compared to forepaw draining brachial LNs. While being consistent with differential expression of costimulatory molecules by migratory cDC2s, and with otherwise limited differences in the number and composition of antigen-bearing migratory cDCs across the sites, it was still possible that insufficient delivery of antigen to brachial LNs after paw immunization was mediating the reduced T cell activation and Th2 programming in this compartment. To test this possibility, we titrated the amount of OVA administered into the skin sites along with a fixed concentration of papain. Although increasing the OVA dose at each site up to 10x the original amount did increase OT-II clonal expansion in both draining LNs, this did not enhance Th2 responses in brachial LNs, suggesting that the total amount of antigen was not a limiting factor in driving reduced Th2 differentiation after footpad immunization (Figure 4E). Of note, increased antigen delivery also did not elicit increased Tbet expression in responding T cells, indicating that high antigen dose availability does not necessarily promote Th1 skewing in papain immunization settings (Figure 4E).

Based on the above observations, we next directly tested the requirements for costimulatory molecule expression on Th2 differentiation *in vivo*. For this, we performed a timed blockade of CD28 by administering an anti-CD28 antibody 24 hours post immunization and harvesting LNs 24 hours later. Delayed anti-CD28 administration allows the T cells to mount initial cognate interactions with cDCs for early priming and activation, but would limit the prolonged costimulatory contacts during the differentiation phase^5^. Indeed, delayed CD28 blockade resulted in very modest reductions of OT-II cellularity, indicating relatively normal initial activation (Figure 5A), but markedly decreased the expression of 4get-GFP, Gata3, BATF, and pS6 in both auricular and brachial LNs (Figure 5A-C). Delayed CD28 blockade also resulted in reduced macro-cluster formation within auricular LNs, instead driving more homogeneous and non-clustered distribution of OT-II cells throughout the T zone, akin to responses observed in footpad draining brachial LNs (Figure 5B, D). Given that costimulation is thought to promote general T cell activation for all helper cell lineages, we next tested whether prolonged costimulatory sensing was important in Th1 inducing settings after CpG OVA immunization. Delayed CD28 blockade elicited much more modest effects on CXCR3 and Tbet expression (Figure S5A), overall indicating that prolonged costimulation was less essential in Th1-inducing settings.

**Figure 5.**
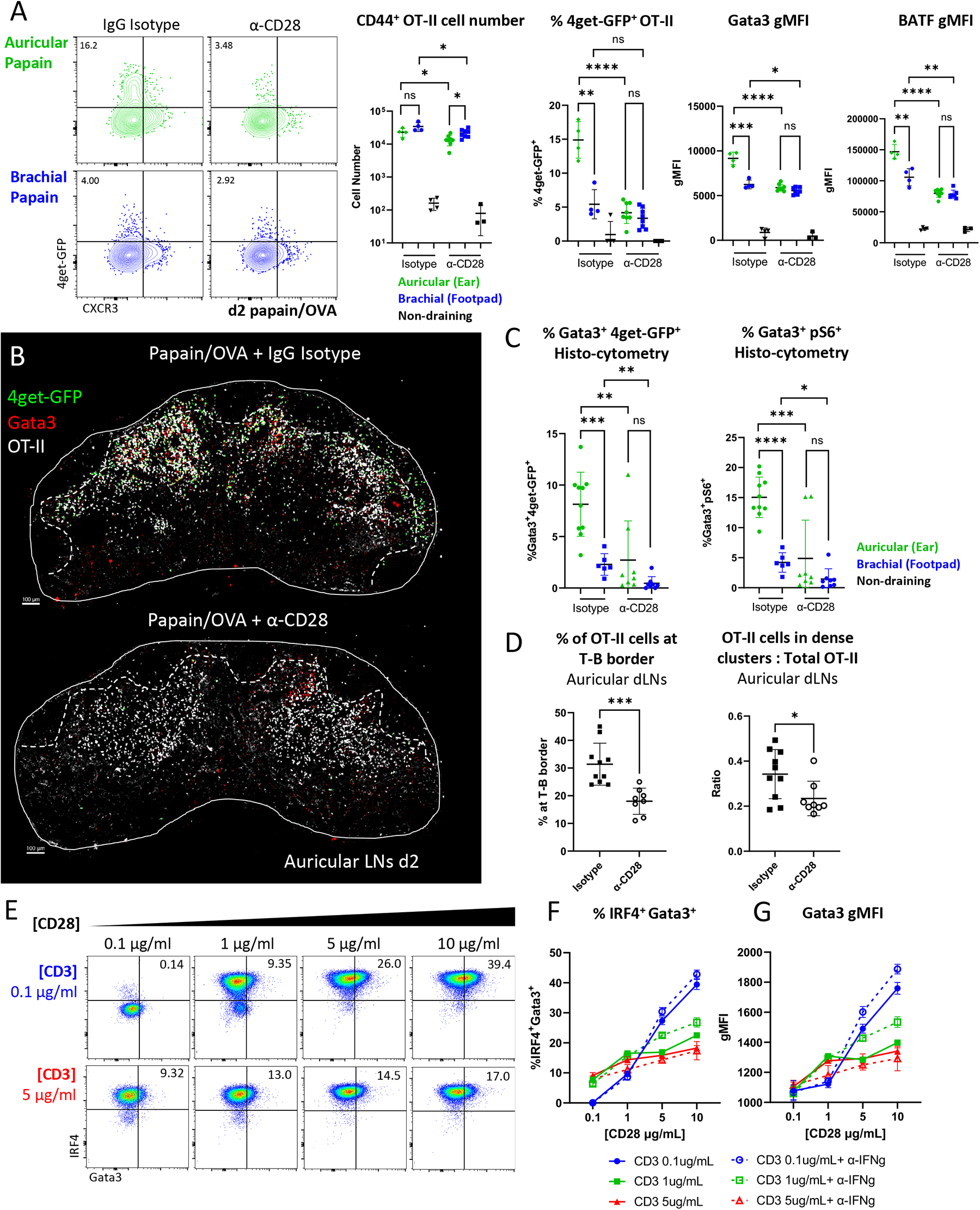
Site-specific T cell responses are mediated through non-equivalent expression of costimulatory molecules by migratory cDCs. A-D) Naïve Ly5.2 4get.OT-II cells were transferred to Ly5.1 mice and injected with papain OVA in the ear pinnae or front footpad. Mice were treated intraperitoneally with α-CD62L 6 hours post immunization, treated intraperitoneally with α-CD28 blocking antibody or IgG isotype control 24 hours post immunization, and dLNs were harvested and assessed by histo-cytometry or flow cytometry at 48 hours. A) Representative flow plots and quantification of CD44^+^ OT-II cell number, 4get-GFP^+^ frequency and Gata3 and IRF4 gMFI of CD44^+^ OT-II cells after α-CD28 or isotype control treatment. B) Representative images depicting OT-II responses in auricular dLN treated with isotype control (top) or α-CD28 blocking antibody (bottom). Dashed lines represent the T-B border. C-D) Histo-cytometry analysis of 4get-GFP and Gata3 expression, OT-II localization, and frequency of macro-clustered OT-II cells. E-G) Naïve OT-II cells were cultured *in vitro* with the indicated concentrations of α-CD3 and α-CD28. α-IFNγ was added to cultures in some conditions (dashed lines). Cells were harvested 48 hours later and assessed for expression of Gata3 and IRF4 by flow cytometry. Representative plots of IRF4 and Gata3 expression are shown with quantification. A-D are representative of at least 2 independent experiments. E-G are representative of 4 independent experiments.

To examine if increasing costimulation can directly modulate Gata3 expression and Th2 differentiation *in vitro*, we next cultured naïve OT-II T cells in non-polarizing culture conditions with varying concentrations of plate-bound anti-CD3 and anti-CD28 for 48 hours. We found that increasing the concentration of available CD28 significantly enhanced Gata3 expression in T cells, and this was particularly evident at low anti-CD3 concentrations (Figure 5E-G). IRF4 was also increased in a CD28-dependent manner, indicating that this transcription factor can be regulated by both TCR and costimulation (Figure 5E-F). Additional blockade of IFN-γ further increased Gata3 expression across both low and intermediate anti-CD3 culture conditions, supporting its role in suppressing Th2 cell differentiation (Figure 5F-G). Moreover, dose-dependent CD28-mediated effects on Gata3 expression were also observed in Th2-polarizing culture conditions, albeit Gata3 and IRF4 expression was significantly greater than in non-polarizing settings (Figure S5B). Together, these data suggest that costimulation can directly enhance T cell activation and Th2 differentiation by inducing IRF4 and Gata3 expression, and that differences in T cell differentiation across the distinct skin draining LNs are most likely driven by non-equivalent expression of costimulatory molecules by migratory cDC2s.

### Th2 macro-clustering and differentiation is driven by prolonged ICAM1-mediated DC-T cell contacts

The above findings indicated that T cell macro-clustering was dependent on prolonged costimulatory molecule availability from migratory cDC2s (Figure 5B). We next tested how prolonged T cell – DC contacts were mediated *in vivo*. The LFA-1 integrin is rapidly upregulated on T cells following activation, and its ligand, ICAM-1, is expressed on DCs after maturation to mediate T cell – DC interactions^46, 76^. LFA-1 has also been shown to reduce the threshold of TCR / costimulatory signaling required for T cell activation thus enabling increased activation in setting of low pMHC-II^77^. We observed marked ICAM-1 upregulation on activated antigen-bearing EαGFP^+^ migratory DCs in both brachial and auricular LNs, and this was also elevated as compared to ICAM-1 expression after CpG immunization (Figure 6A). To test if T cell macro-clustering and Th2 differentiation was mediated via prolonged integrin-driven T – DC interactions, we performed a delayed LFA-1 blockade, treating mice with an anti-LFA-1 blocking antibody 24 hours post papain OVA administration and examining responses in auricular and brachial LNs 24 hours later^46, 78^. Delayed LFA-1 blockade did not markedly alter OT-II T cell proliferation, indicating relatively normal initial activation (Figure 6C). In contrast, 4get-GFP, Gata3, and IRF4 expression were all significantly decreased in draining LNs after delayed LFA-1 blockade, indicating reduced Th2 differentiation (Figure 6B-C, E). LFA-1 blockade also disrupted the T cell macro-clustering at the T-B border, indicating limited formation of Th2 microenvironments (Figure 6D, F). In contrast to type-II inflammatory settings, delayed LFA-1 blockade had a minimal effect on Th1 differentiation after OVA CpG immunization (Figure S6A-B), suggesting that prolonged LFA-1 integrin mediated cell interactions are less important in these settings.

**Figure 6.**
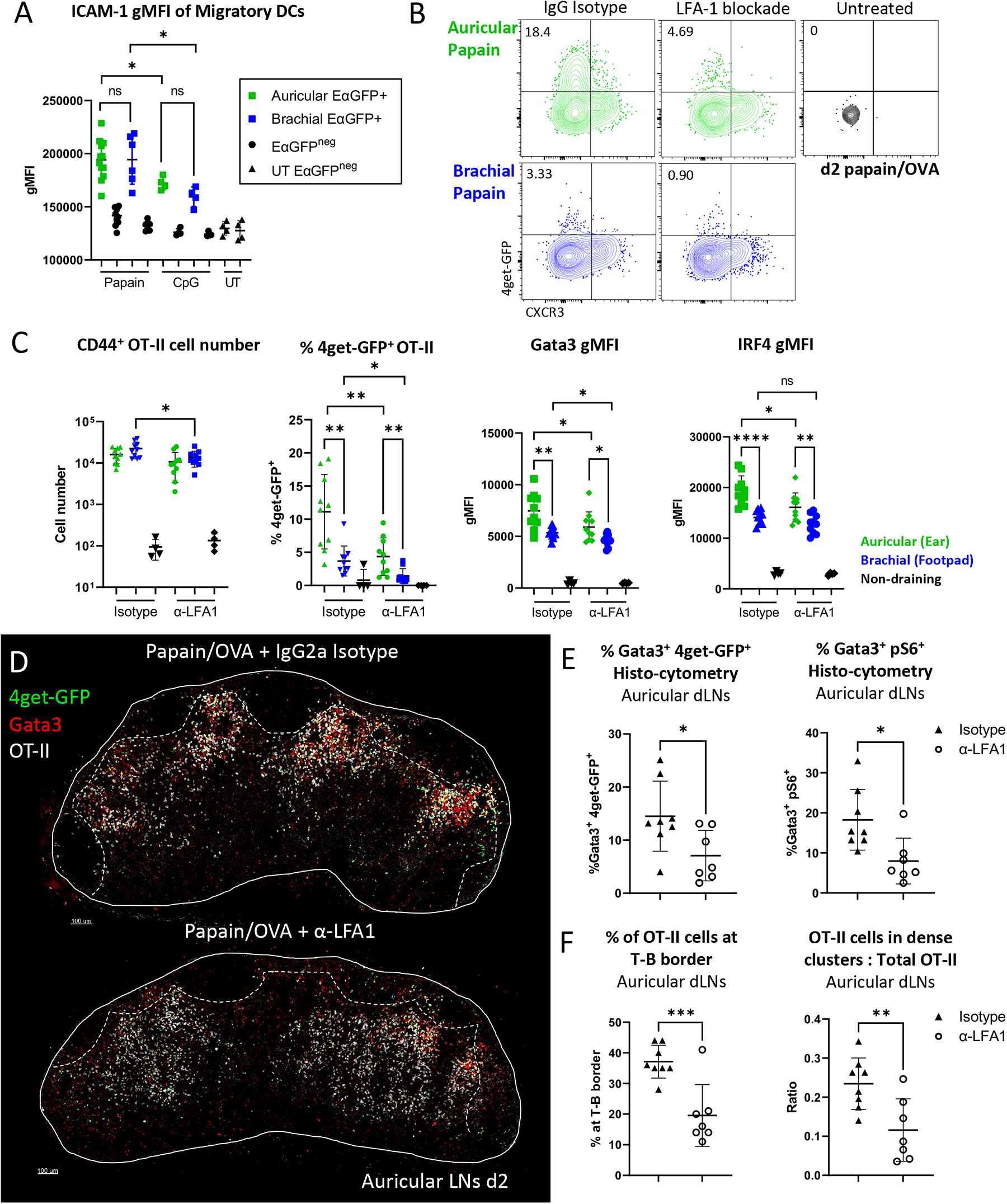
Th2 macro-clustering and differentiation is driven by prolonged ICAM-1 mediated DC-T cell contacts. A) B6 mice were immunized in the ear pinnae and front footpad with papain or CpG plus EαGFP and harvested for flow cytometry 48 hours later. Expression of ICAM-1 on EαGFP^+^ and EαGFP^neg^ or naïve migratory DCs in the indicated LNs is shown. B-F) Naïve Ly5.2 4get.OT-II cells were transferred to Ly5.1 mice and injected with papain OVA in the ear pinnae or front footpad. Mice were treated intraperitoneally with α-CD62L 6 hours post immunization, treated intraperitoneally with α-LFA1 blocking antibody or IgG isotype control 24 hours post immunization, and dLNs were harvested and assessed by flow cytometry and imaging at 48 hours. B) Representative flow plots showing 4get-GFP and CXCR3 expression on CD44^+^ OT-II cells. C) Quantification of the number of CD44^+^ OT-II cells, frequency of 4get-GFP^+^ cells, and Gata3 and IRF4 gMFI of CD44^+^ OT-II cells after α-LFA1 or isotype control treatment. D) Representative microscopy images depicting OT-II responses in auricular dLN treated with isotype control (top) or α-LFA1 blocking antibody (bottom). Dashed lines represent T-B border. E-F) Histo-cytometry analysis of 4get-GFP and Gata3 expression, OT-II localization, and frequency of macro-clustered OT-II cells in auricular LNs with the indicated treatments. A, D-F are representative of 2 independent experiments. B-C are representative of 4 independent experiments.

After activation, T cells can also express ICAM-1 on the cell surface, and in addition to T – DC interactions could potentially engage in homotypic T – T cell contacts^46^. To dissect whether delayed LFA-1 blockade was primarily affecting T – DC or T – T cell interactions, we generated ICAM1.KO OT-II cells (Figure S6C) and compared these responses to WT OT-II cells. In these experimental settings, ICAM1.KO OT-II T cells retain normal LFA-1 integrin functionality and can interact with ICAM1-expressing cDCs but lack the ability to engage in homotypic T – T cell contacts. We found no defects in Th2 differentiation in ICAM-1 deficient OT-II T cells, indicating that LFA-1 / ICAM-1 mediated homotypic T – T interactions are not required for early Th2 responses (Figure S6D). Together, these data suggest that enhanced T cell activation via costimulatory molecules expressed by cDCs promotes prolonged LFA1 integrin-mediated T – DC interactions within auricular ear draining LNs, and these contacts in turn drive the formation of T cell macro-clusters and promote Th2 differentiation.

### Increased costimulation in auricular draining LNs promotes enhanced cytokine signaling and Th2 differentiation

Costimulation promotes the production of IL-2 by activated T cells, as well as can elicit the production of the Th2 cytokine IL-4 *in vitro*^4, 7–11, 65, 66^. IL-2 signaling in turn increases expression of the IL-4 receptor, IL-4Rα, such that IL-2 stimulated T cells become more receptive to IL-4^79^. T cell costimulation and macro-clustering may thus promote increased local bioavailability of the cytokines IL-2 and IL-4, produced directly by activated T cells, which may reinforce localized Th2 differentiation within the macro-clusters. Indeed, we found CD25 was highly expressed on the activated CD44^+^ Gata3^+^ OT-II Th2 cells (Figure 7A). Further, early Gata3^+^CD25^+^ Th2 cells had increased expression of phosphorylated (p)STAT5 and pSTAT6, indicating enhanced and/or sustained IL-2 and IL-4 signaling, respectively, as compared with Gata3^neg^CD25^neg^ non-Th2 cells (Figure 7B).

**Figure 7.**
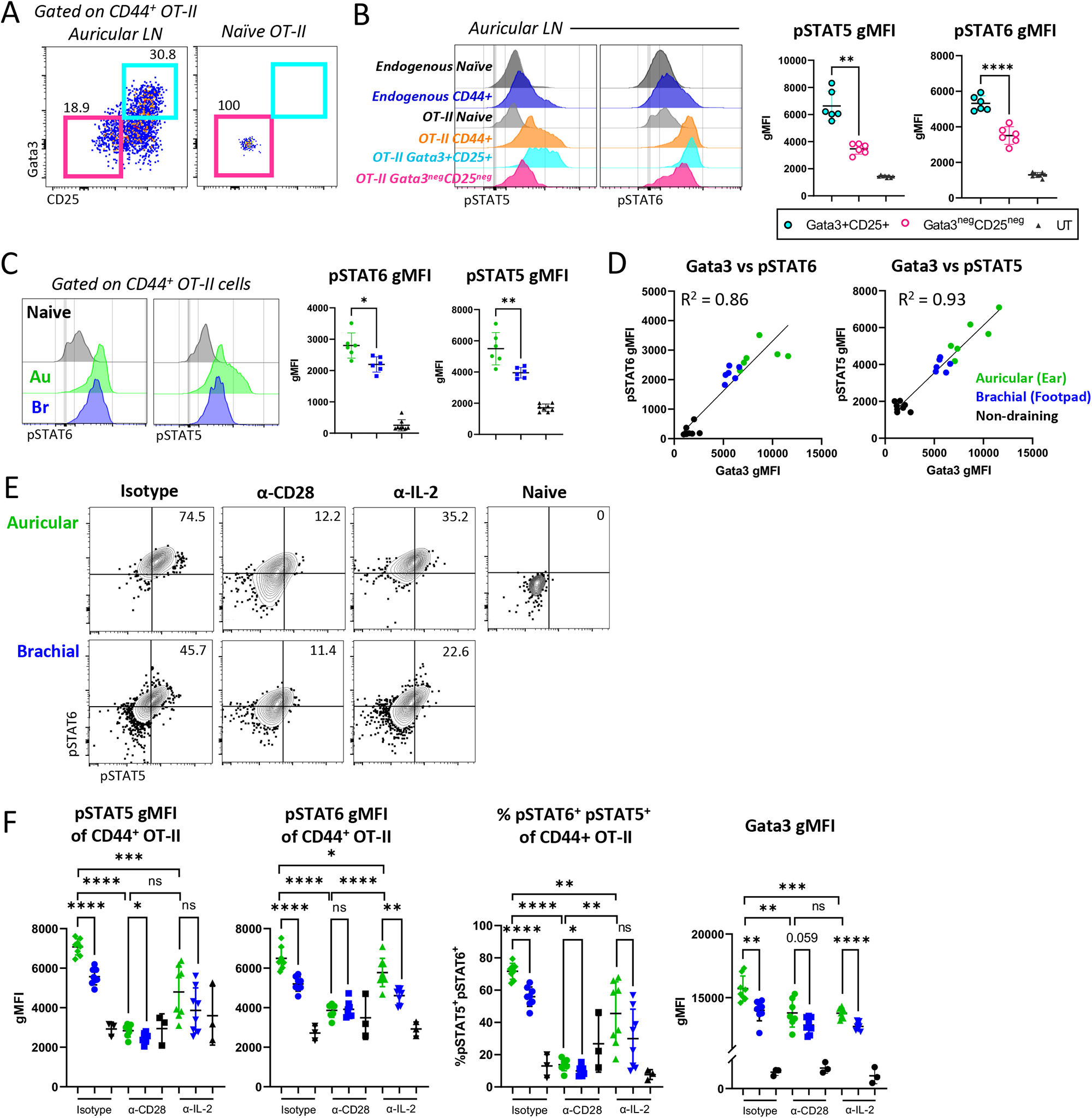
Increased costimulation in auricular draining LNs promotes increased cytokine signaling and Th2 differentiation. A-D) Naïve Ly5.1 OT-II T cells were transferred to B6 mice and injected with papain OVA in the ear pinnae or front footpad. Mice were treated intraperitoneally with α-CD62L blocking antibody 6 hours post immunization and dLNs were harvested and assessed by flow cytometry at 48 hours. A) Representative flow plots of Gata3 and CD25 expression on CD44^+^ OT-II cells in auricular dLNs. B) Representative histograms and quantification of pSTAT5 and pSTAT6 staining for the indicated cell subsets within the same sample. C) Representative histograms and quantification of pSTAT5 and pSTAT6 expression in auricular and brachial LNs and D) pSTAT5 and pSTAT6 correlation with Gata3 gMFI on CD44^+^ OT-II cells is shown. E-F) Naïve Ly5.1 OT-II cells were transferred to B6 mice and injected with papain OVA in the ear pinnae or front footpad. Mice were treated intraperitoneally with α-CD62L 6 hours post immunization, treated intraperitoneally with α-CD28 blocking antibody, α-IL-2 blocking antibody, or IgG isotype control 24 hours post immunization, and dLNs were harvested and assessed by flow cytometry at 48 hours. Quantification of frequency of pSTAT5^+^ and pSTAT6^+^ cells, gMFI of pSTAT5 and pSTAT6, and Gata3 expression on CD44^+^ OT-II cells are shown. Data are representative of at least 2 independent experiments.

Consistent with non-equivalent T cell activation and differentiation in different draining LNs, both pSTAT5 and pSTAT6 in responding OT-II cells were significantly elevated in auricular as compared to brachial draining LNs (Figure 7C). pSTAT levels were also directly correlated with non-equivalent GATA3 expression across the sites (Figure 7D), suggesting direct involvement in Th2 differentiation. We thus tested whether differential sensing of costimulatory molecules by the responding T cells in distinct draining LNs resulted in non-equivalent cytokine signaling and Th2 differentiation. For this, we again utilized the timed CD28 blockade and examined phosphorylation of STAT5 and STAT6 molecules 24 hours later. Indeed, we found that delayed anti-CD28 treatment significantly reduced pSTAT5 and pSTAT6 expression in auricular draining LNs, and completely abrogated the differences between the tissues (Figure 7E-F). Similar effects were observed after *in vivo* anti-IL-2 blockade, and both anti-CD28 and anti-IL-2 treatments resulted in comparable decreases in STAT5, and to a lesser extent STAT6, phosphorylation (Figure 7E-F). Finally, both anti-CD28 and IL-2 blockade markedly reduced Gata3 expression in responding OT-II cells indicating a clear link between prolonged costimulation, cytokine sensing, and Th2 differentiation (Figure 7F). In sum, our data indicate that increased sensing of costimulatory molecules by early differentiating T cells in ear draining auricular LNs is associated with enhanced T cell macro-clustering, increased local IL-2 and IL-4 cytokine production and sensing, and amplified early Th2 differentiation (Figure S7).

## Discussion

*In vivo* mechanisms driving early Th2 differentiation in LNs have remained enigmatic, in large part due to the complex role of TCR signaling in directing T cell effector fates and the lack of clear understanding of which molecules expressed by cDCs, or other innate cells in LNs, selectively induce Th2 polarization^21^. Our studies indicate that early differentiating Th2 cells undergo enhanced T cell signaling and activation, and that this is mediated through macro-clustering at the T-B border with migratory cDC2s displaying high levels of costimulatory molecules and integrin ligands but relatively low levels of surface pMHC. This in turn appears to promote efficient cytokine exchange, in particular for IL-2 and IL-4, among neighboring activated T cells to reinforce localized Th2 differentiation and to support prolonged proliferation. Thus, formation of discrete spatial microenvironments within LNs in which T cells integrate quantitatively strong activation signals from cDC2s coupled with qualitative cytokine sensing from nearby T cells promotes the initiation of large-scale Th2 responses *in vivo*.

Past studies have demonstrated that *in vitro* stimulation of CD4 T cells with strong TCR agonists or high dose of peptide promotes Th1 differentiation, while low dose signals elicit Th2 differentiation^2, 3^. This has also been supported by *in vivo* work using peptide-pulsed DCs, which showed that TCR signal strength serves as a rheostat to control cytokine receptor expression, in turn modulating the ability of T cells to sense cytokines and undergo T helper cell differentiation^2–4^. Our studies are consistent with this hierarchy model, showing that cells undergoing the greatest degree of activation also receive abundant cytokines and thus become more differentiated. However, we show that during *in vivo* allergen exposure, prolonged and enhanced T activation leads to Th2 and Tfh, not Th1 polarization, and that Th1 differentiation occurs at minimal rates for these responses regardless of antigen dosage. Differences in these observations are likely explained by the fact that environmental allergens such as papain elicit inflammatory functions via proteolytic disruption of the epithelial barrier and release of the alarmin IL-33^80^, and this in turn induces cDC maturation and costimulatory molecule expression through local type 2 innate lymphoid cell (ILC2) activation^13, 14, 81, 82^, thus necessitating *in vivo* administration to appropriately instruct cDCs. In addition, we and others show that during type-II inflammation, cDCs have a reduced capacity to produce the type-I skewing cytokine IL-12, thus minimizing the capacity of T cells to undergo Th1 differentiation^34, 38^. Further, activated T cells displaying high levels of cytokine receptors appear confined within microenvironments rich in other T cells and cDC2s and away from other potential cellular sources of IL-12, which could suppress early Th2 differentiation^35, 36^. In this regard, we previously showed that papain administration does not elicit robust monocyte recruitment to the draining LNs^38^, as these cells can also produce copious amounts of IL-12 during type-I inflammation^83^. Altogether, these findings suggest there is limited availability of Th1 inducing factors in draining LNs after papain administration, indicating that highly activated T cells undergoing prolonged interactions with cDCs receive Th2 polarizing cytokines in the absence of Th1-promoting factors, leading to the generation of large-scale Th2 responses. Notably, prolonged costimulatory signaling and formation of T cell macro-clusters appears far less critical for Th1 responses *in vivo*. This may be due to comparatively lower expression of integrin ligands by cDC2s to mediate prolonged clustering, high abundance of inflammatory cytokines across vast regions of the LN parenchyma, and presence of chemokines which would drive CXCR3-expressing early Th1 cells away from sites of initial T – DC contacts^38, 39^.

Extensive evidence now exists that migratory cDC2s are required for Th2 differentiation^25–30^. We similarly find that ablation of cDC2 migration from the skin abrogates Th2, as well as Tfh, differentiation in LNs, albeit not necessarily at the cost of reduced early T cell priming. This may be explained by the fact that in settings of ample antigen drainage, lymph node resident cDCs initiate early T cell proliferation, while migratory cDC2s induce downstream Th2 differentiation^37, 38, 60^. Factors expressed by cDC2s to selectively initiate Th2 skewing have remained unknown^34^, but costimulatory molecule expression by cDCs can promote type-II cytokine production by *in vitro* stimulated T cells^7–12^. *In vivo*, TSLP-driven OX40L costimulatory molecule expression by cDCs has been positively linked with Th2 responses^14, 15, 84^.

Costimulatory molecules on their own do not constitute a Th2 polarizing stimulus and can be involved in promoting general T cell activation. However, we find that prolonged costimulation was less essential for adjuvant induced Th1 responses, indicating differential requirements of costimulation for distinct helper cell lineages. Our data do support the notion that costimulation is essential for inducing IL-2 and IL-4 cytokine production by activated T cells^18–21, 85–87^, and likely for eliciting IL-4Rα upregulation, in turn allowing IL-2-experienced T cells to become more receptive to IL-4 signaling to drive enhanced Th2 differentiation^79^. IL-2 also skews responding T cells away from the Tfh lineage, thus supporting additional specification of helper cell fate^51, 55, 56^. Conversely, the spatial proximity of macro-clusters near B cell follicles could also directly support Tfh differentiation by enhancing the probability of interactions between those activated T cells which receive less IL-2 and neighboring B cells presenting cognate antigens, thereby explaining the involvement of migratory cDC2s in both Th2 and Tfh responses^88^.

The initial cellular source of IL-4 in draining LNs has not been clearly defined, yet recently stimulated T cells can produce IL-4 in a TCR-dependent manner *in vitro*^19, 20^. Mice deficient in IL-4Ra also retain the capacity to secrete IL-4, suggesting that T cells can produce IL-4 without a requirement for prior IL-4 sensing^89^. Consistent with this, IL-4 secretion by T cells can be achieved by the activation of BATF/IRF4/Jun complexes downstream of TCR engagement and costimulation in a Gata3-independent manner^67–75^. Indeed, we observe robust expression of IRF4 and BATF in activated T cells within ear draining auricular LNs, suggesting these transcription factors may be sufficient to induce initial IL-4 production by responding T cells within macro-clusters which can then be further amplified by canonical IL-4/IL-4Rα driven Gata3 expression. While our studies do not examine other potential sources of IL-4 in LNs, the majority of activated OT-II T cells express pSTAT6, indicative of more widespread cytokine sensing after papain administration^90^. Our findings do suggest that T cells within the macro-clusters experience more extensive cytokine signaling than those outside the clusters, and this further enhances Th2 differentiation within local microenvironments. Supporting this are past observations that both IL-2 and IL-4 are secreted by T cells in a broadcasted, non-polarized fashion^91, 92^, as well as our results demonstrating that late CD28 blockade, which disrupts macro-clustering, results in diminished STAT phosphorylation.

Of note, our data also suggest that during papain exposure, antigen-bearing migratory cDC2s have low levels of pMHC complexes on the cell surface, accompanied by high amounts of costimulatory molecules and integrin ligands. Mechanisms of how type-II inflammation impacts antigen processing and presentation, as well as whether it extends to other type-II settings remains to be determined, but could be driven in part by modulating the cytoskeletal properties of cDCs^93^. Together with TCR engagement, costimulation enhances LFA-1 integrin activation, which enables prolonged T cell – cDC interactions, as well as reduces the threshold of TCR signaling required for T cell activation, overall being consistent with our findings that low levels of pMHC-II can still lead to robust T cell responses^77^. Additional integrin-mediated homotypic interactions among activated T cells may also take place, albeit likely not via ICAM-1 / LFA-1 interactions. Of note, a recent study has demonstrated a role for the integrin αVβ3, for Th2 differentiation *in vitro*, and this integrin has also been previously linked with migratory behavior of Th2 cells in inflamed tissues^94, 95^.

A key unexpected finding in our work was that not all types of skin generated equivalent Th2 responses in the draining LNs. Our studies primarily focused on the ear vs footpad draining LNs, but additional variation across the skin is likely. A major unanswered question remains as to why cDCs within different skin sites are non-equivalently activated after exposure to the same agonist and whether this extends to other barrier tissues in mice and humans. Homeostatic tissue signals like IL-13 and IL-18 can impart cDCs and ILC2s with a tissue-specific identity, potentially reflecting intrinsic pre-programming of cDCs and other cells based on skin type residence^96–98^. In addition, differences could reflect divergent cDC experiences at the site of immunization. Many cytokines, including alarmins, are released from epithelial and stromal cells during type-II inflammation and barrier disruption events, and whether these cytokines are equivalently released between skin sites requires further study. There could be many additional tissue specific adaptations, such as neuronal composition, mast cell differences, or divergent microbiomes which direct dermal cDC2 maturation and activation^99–102^. Epidermal barrier thicknesses and other structural differences between skin sites could also contribute^103^. Therefore, skin should not be thought of as a single barrier tissue, but unique compartments which may respond differently to environmental triggers. A possible evolutionally benefit of reduced Th2 responses after footpad allergen administration could reflect the need for dampened inflammation in tissues with constant mechanical stress, environmental exposure, and likely damage, while still retaining the ability to induce robust Th1 responses to microbial challenges. Finally, a common feature of Th2 allergic responses is atopic march development, in which initial skin sensitization leads to downstream pathology across peripheral organs. It will thus be important to understand how different regions of the skin, the largest barrier tissue in the body, respond to allergen exposure and drive disease development.

## Figure Legends

**Figure S1.**
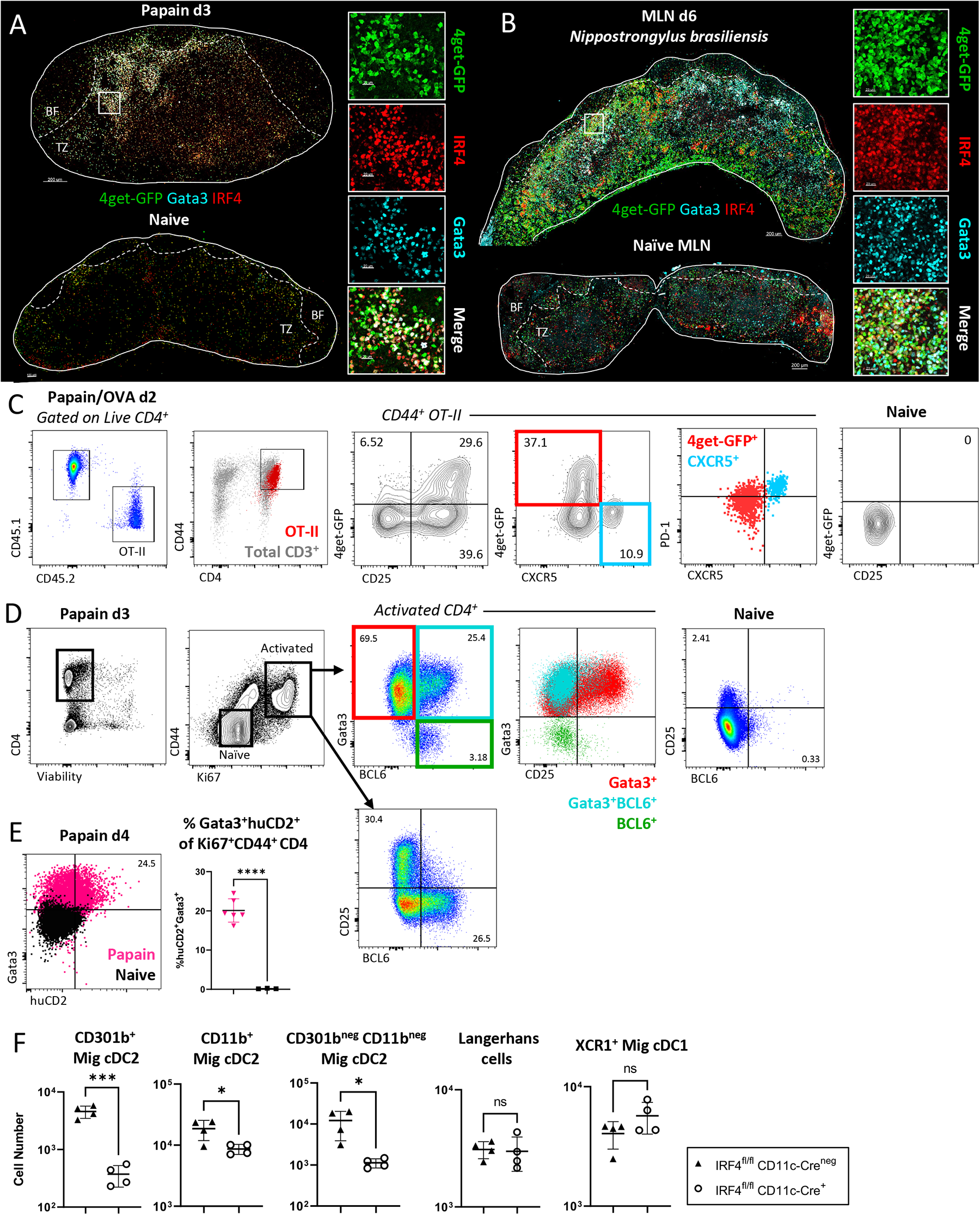
Cellular composition of Th2 microenvironments after papain immunization or *N.b.* infection. A) 4get-GFP mice were immunized in the ear pinnae with papain and auricular dLNs or non-draining naïve LNs were harvested for confocal microscopy 3 days later. Representative images and zoom ins are shown with the indicated markers. B) 4get-GFP mice were inoculated subcutaneously at the tail base with 500 *N.b.* L3 larvae and the draining mesenteric LN (MLN) or MLN from naïve mice were harvested 6 days later for confocal microscopy. Representative image and zoom in depicting Th2 macro-clustering in the MLN with the indicated markers is shown. C) Naïve Ly5.2 4get.OT-II T cells were transferred to Ly5.1 mice and injected with papain OVA in the ear pinnae, treated with α-CD62L blocking antibody at 6 hours, and harvested for flow cytometry 48 hours later. Representative gating for T follicular helper (Tfh) and T effector (Teff) OT-II cells is shown with the indicated markers. D) B6 mice were immunized with papain in the ear pinnae and auricular dLNs were harvested 3 days later for flow cytometry. Representative gating for endogenous T cell activation is shown with Gata3, BCL6, and CD25 identifying Tfh and Teff T cells. E) KN2^+/-^ mice were immunized with papain in the ear pinnae and auricular dLNs or naïve lymph nodes (nLN) were harvested 4 days later for flow cytometry. Representative plot and quantification of huCD2 and Gata3 expression is shown. F) CD11c-Cre^+^ IRF4^fl/fl^ or CD11c-Cre^neg^ IRF4^fl/fl^ were immunized with papain in the ear pinnae and dLNs were harvested 2 days later for flow cytometry. Quantification of the indicated myeloid cell subsets is shown. BF = B cell follicles; TZ = T cell zone. Dashed lines represent T-B border. A, C-D are representative of 5 independent experiments. B, E-F are representative of 2-3 independent experiments.

**Figure S2.**
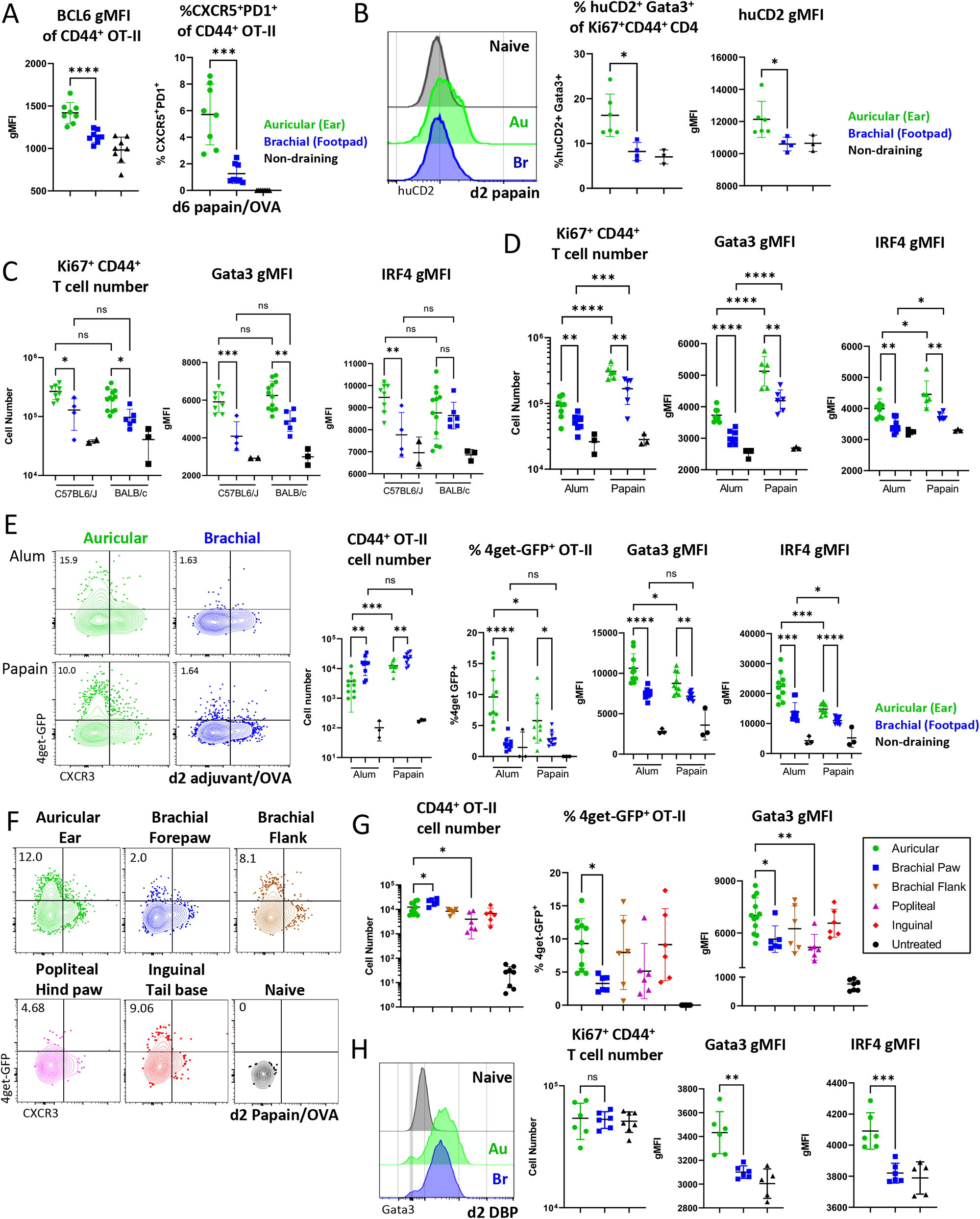
Site-specific Th2 responses are maintained across mouse strains and adjuvants. A) 0.5×10^6^ naïve Ly5.2 4get.OT-II cells were transferred to Ly5.1 mice and injected with papain OVA in the ear pinnae or front footpad and dLNs, spleen, and lung were harvested and assessed by flow cytometry 6 days later. BCL6 gMFI of CD44^+^ OT-II cells and frequency of CXCR5^+^PD-1^+^ CD44^+^ OT-II cells are shown. B) KN2^+/-^ mice were immunized with papain in the ear pinnae or front footpad and dLNs were assessed by flow cytometry 2 days later. Representative plot and frequency of KN2^+^Gata3^+^ and KN2 gMFI of CD44^+^Ki67^+^ CD4 cells are shown. C) B6 or Balb/c mice were immunized in the ear pinnae and front footpad with papain and dLNs were harvested at day 3 for flow cytometry. Activated CD44^+^Ki67^+^ T cell number, and Gata3 and IRF4 gMFI of CD44^+^Ki67^+^ T cells are shown. D) B6 mice were immunized in the ear pinnae and front footpad with papain or Alum OVA and dLNs were harvested at day 3 for flow cytometry. Activated CD44^+^Ki67^+^ T cell number, and Gata3 and IRF4 gMFI of CD44^+^Ki67^+^ T cells are shown. E) Naïve Ly5.2 4get.OT-II cells were transferred to Ly5.1 mice, injected with papain OVA or Alum OVA in the ear pinnae or front footpad, treated intraperitoneally with α-CD62L 6 hours post immunization and dLNs were harvested and assessed by flow cytometry at 48 hours. Representative plots, CD44^+^ OT-II cell number, expression of 4get-GFP of CD44^+^ OT-II cells, and Gata3 and IRF4 gMFI of CD44^+^ OT-II cells are shown. F-G) Naïve Ly5.2 4get.OT-II cells were transferred to Ly5.1 mice, and injected with papain OVA in the ear pinnae, front footpad, hind footpad, dorsal flank skin, or tail base. Mice were treated intraperitoneally with α-CD62L 6 hours post immunization and dLNs were harvested and assessed by flow cytometry at 48 hours. Representative plots, CD44^+^ OT-II cell number, frequency of 4get-GFP^+^ cells and Gata3 gMFI of CD44^+^ OT-II cells are shown. H) DBP was topically applied (painted) onto the ear or front footpad skin of mice. dLNs were harvested for flow cytometry 48 hours later. Activated CD44^+^Ki67^+^ T cell number, and Gata3 and IRF4 gMFI of CD44^+^Ki67^+^ T cells are shown. A-G are representative of 2-3 independent experiments. H is representative of 4 independent experiments.

**Figure S3.**
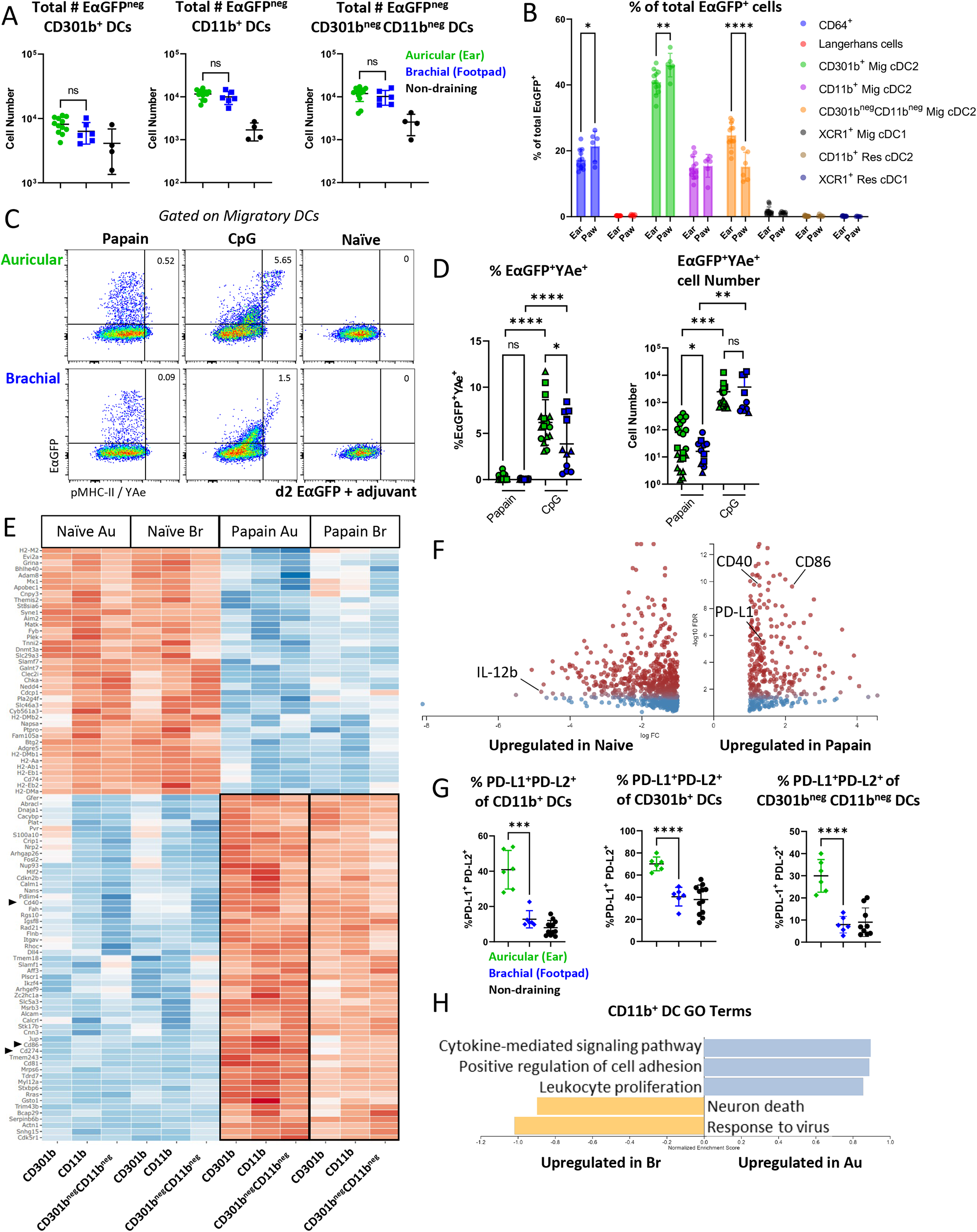
Transcriptional signatures of naïve and activated cDC2s. A-D) B6 mice were immunized in the ear pinnae or front footpad with papain or CpG and EαGFP as indicated and harvested 48 hours later. A) Total number of EαGFP^neg^ cells for the indicated DC subsets. B) Frequency of EαGFP^+^ cells for the indicated cell subsets. C-D) Representative flow plots and quantification showing EαGFP and YAe expression on migratory DCs after immunization with the indicated adjuvant. Symbol shape represents individual experiments (three total). E-F) EαGFP^+^ DC populations were sorted from dLNs on day 2 for bulk RNA sequencing. E) Heatmap of all DEGs with a FDR<0.05 and greater than 2x fold change between naïve and papain immunized DCs from auricular and brachial LNs is shown. F) Volcano plot of DEGs between papain immunized and naïve DCs from auricular and brachial dLNs. Red indicates a FDR<0.05. Genes with a log fold change greater than 2 are shown. G) Migratory DCs were assessed for surface PD-L1 and PD-L2 expression by flow cytometry. Quantification of the frequency of PD-L1 and PD-L2 for the indicated DC subsets is shown. H) GO term enrichment analysis based on all DEGs in EαGFP^+^ CD11b^+^ DC populations between auricular and brachial dLNs. Au = auricular; Br = brachial. A-D, G are representative of 3 independent experiments. E-F and H are representative of 1 independent RNA sequencing experiment with n=3 per group.

**Figure S5.**
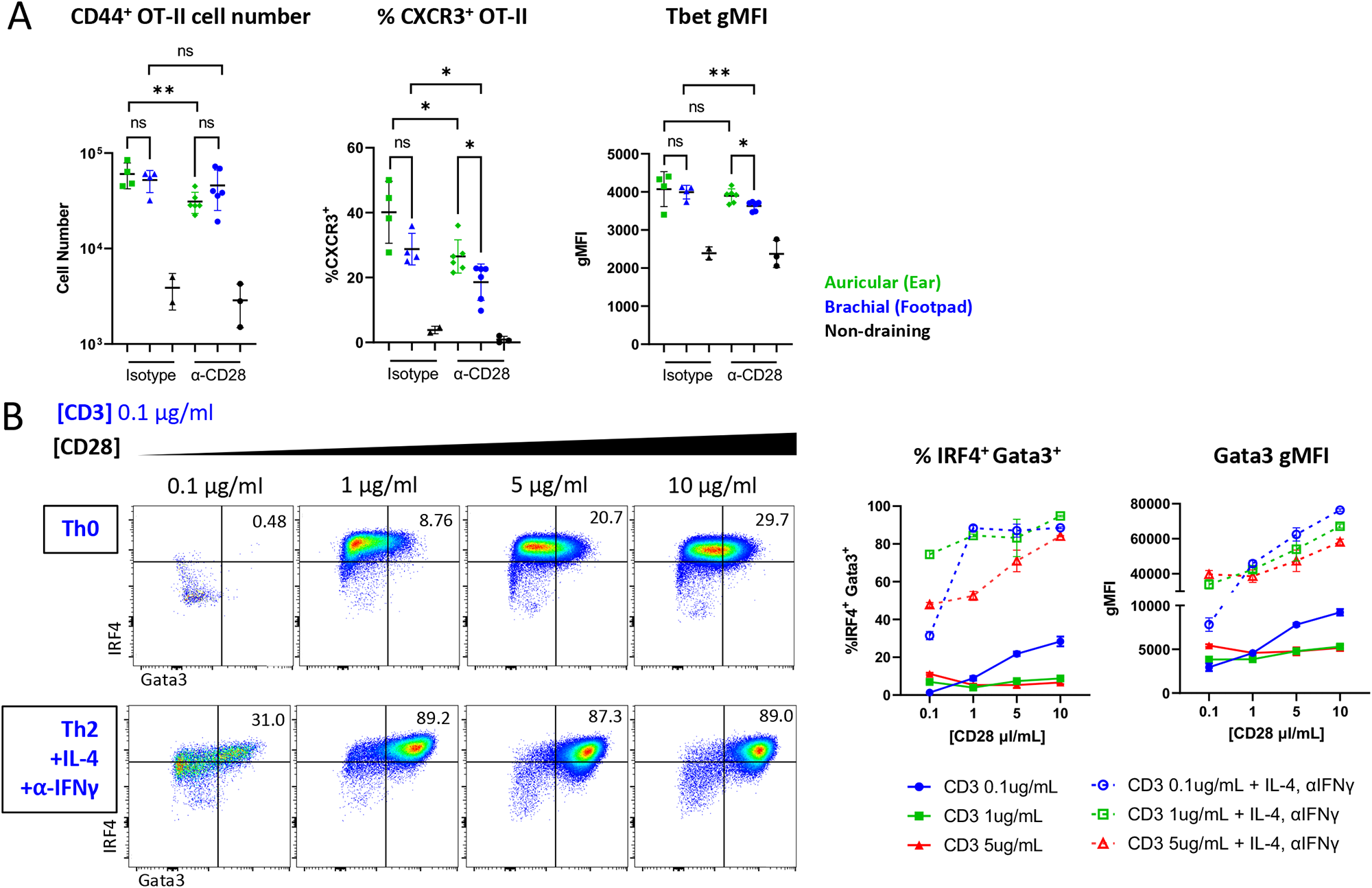
Late CD28 blockade has a minimal effect on Th1 differentiation. A) Naïve Ly5.2 4get.OT-II cells were transferred to Ly5.1 mice and injected with CpG OVA in the ear pinnae or front footpad. Mice were treated intraperitoneally with α-CD62L 6 hours post immunization, treated intraperitoneally with α-CD28 blocking antibody or IgG isotype control 24 hours post immunization, and dLNs were harvested and assessed by flow cytometry at 48 hours. Quantification of CD44^+^ OT-II cell number, frequency of CXCR3^+^ cells, and Tbet gMFI on CD44^+^ OT-II cells are shown. B) Naïve OT-II cells were cultured *in vitro* with the indicated concentrations of α-CD3 and α-CD28. IL-4 and α-IFNγ were added to the culture in some conditions (dashed lines). Cells were harvested for flow cytometry 48 hours later and assessed for expression of Gata3 and IRF4. Representative plots of IRF4 and Gata3 expression are shown with quantification. A is representative of 2 independent experiments. B is representative of 4 independent experiments.

**Figure S6.**
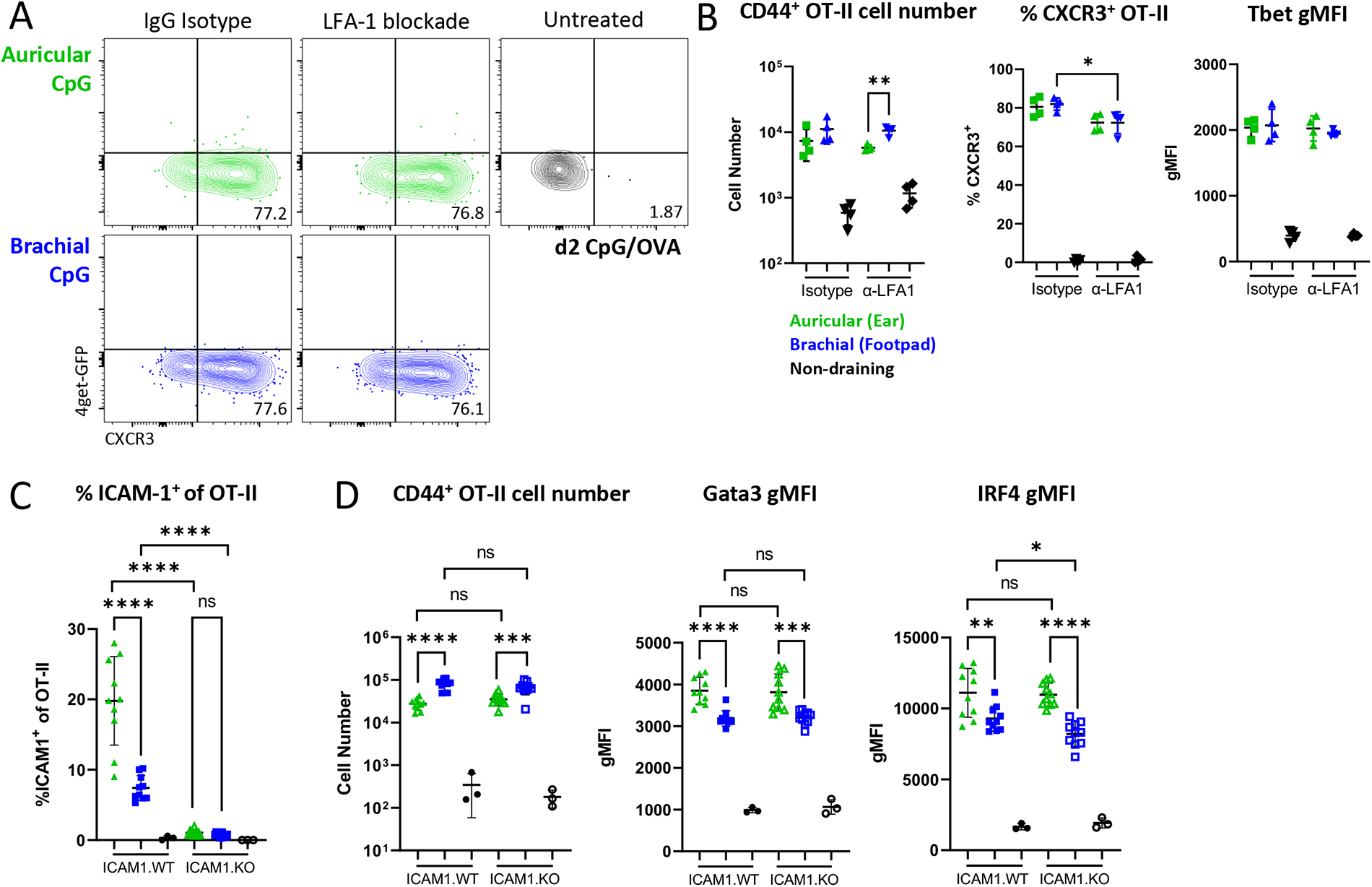
ICAM-1 expression on T cells is dispensable for Th2 differentiation. A-B) Naïve Ly5.2 4get.OT-II cells were transferred to Ly5.1 mice and injected with CpG OVA in the ear pinnae or front footpad. Mice were treated intraperitoneally with α-CD62L 6 hours post immunization, treated intraperitoneally with α-LFA1 blocking antibody or IgG isotype control 24 hours post immunization, and dLNs were harvested and assessed by flow cytometry at 48 hours. A) Representative plots showing 4get-GFP and CXCR3 expression on CD44^+^ OT-II cells. B) Quantification of CD44^+^ OT-II cell number, frequency of CXCR3^+^ cells, and Tbet gMFI on CD44^+^ OT-II is shown. C-D) Naïve ICAM-1.KO or ICAM-1.WT OT-II cells were transferred to CD45.1^+^ B6 recipients and injected with papain OVA in the ear pinnae or front footpad, treated intraperitoneally with α-CD62L 6 hours post immunization, and dLNs were harvested and assessed by flow cytometry at 48 hours. C) Frequency of ICAM-1^+^ OT-II cells from ICAM-1.WT and ICAM-1.KO mice. D) Quantification of the number of CD44^+^ OT-II cells, and Gata3 and IRF4 gMFI of CD44^+^ OT-II cells. WT = wild type; KO = knockout. Data are representative of 2 independent experiments.

**Figure S7.**
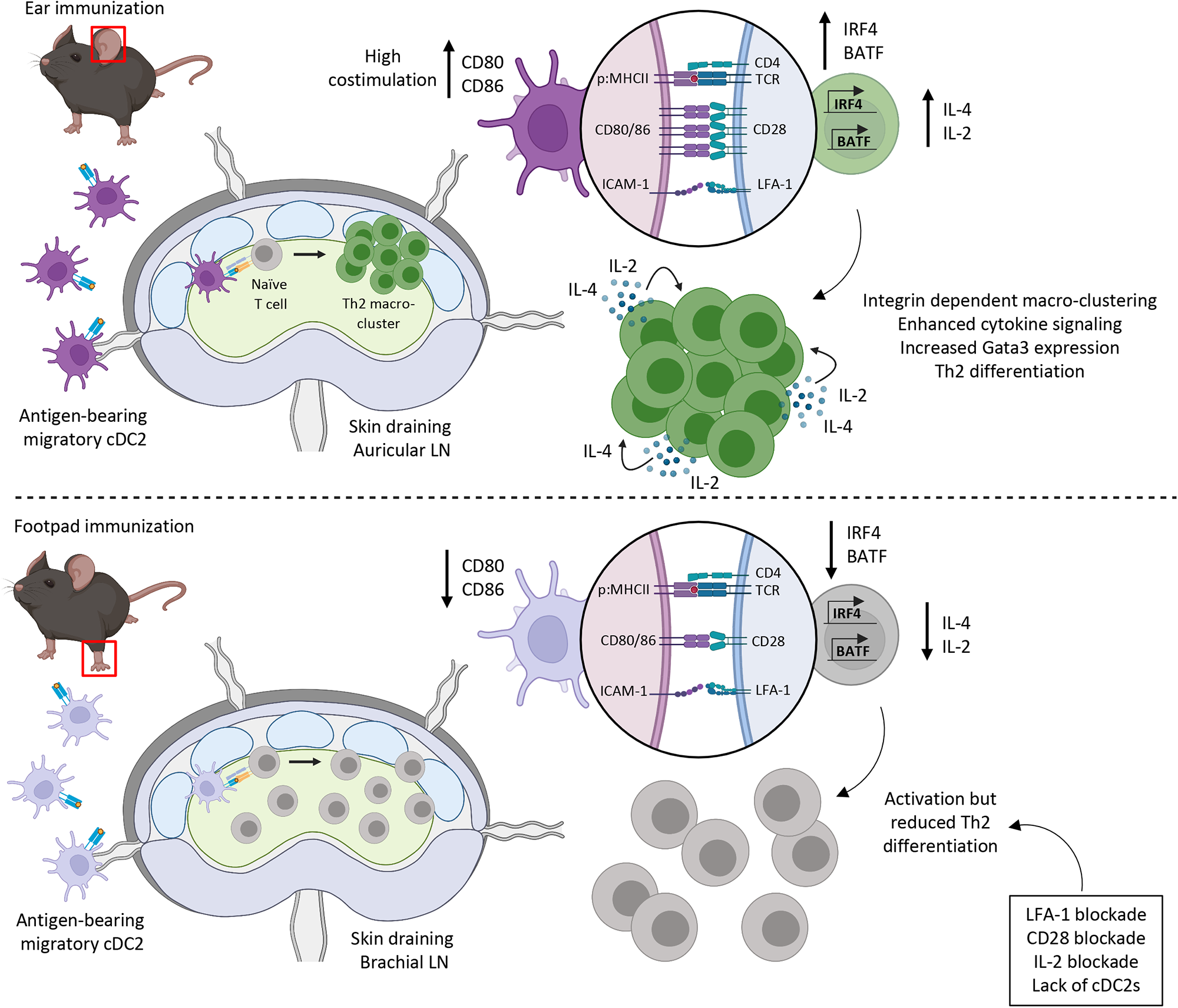
Proposed model for initiation of Th2 differentiation in skin draining LNs. Papain immunization of the skin elicits the maturation and migration of migratory antigen-bearing cDC2 to draining LNs where they induce Th2 responses within dedicated microenvironments localized at the T-B border. Th2 differentiation is driven by prolonged T – DC contacts leading to T cell activation through low levels of pMHC and high levels of costimulatory molecules on cDC2s thus driving integrin-mediated macro-clustering, efficient cytokine exchange, and localized Th2 differentiation. Certain skin sites, such as the paw, induce reduced expression of costimulatory molecules on cDC2s which leads to reduced Th2 responses within corresponding draining LNs.

## Materials & Methods

### Mice

C57BL/6J, BALB/cJ, B6.Cg-Tg(Itgax-cre)1-1Reiz/J (CD11c-Cre), B6.129S1-Irf4^tm1Rdf^/J (IRF4^fl/fl^), and B6.129S4-*Icam1^tm1Jcgr^*/J (ICAM-1.KO) were obtained from Jackson Laboratory. B6.SJL-*Ptprc^a^Pepc^b^*/BoyCrl (CD45.1) mouse stain was obtained from Charles River Laboratory. CD45.1^+^ B6.Cg-Tg(TcraTcrb)425Cbn/J (OT-II) mice were obtained from a donating investigator (P.J. Fink, University of Washington) and crossed with CD45.2^+^ C57BL/6J mice to generate CD45.1^+^ OT-II and CD45.1^+^CD45.2^+^ OT-II lines that were used interchangeably based on recipient congenic status. The KN2 and 4get-GFP mouse strains on a B6 background were obtained from a donating investigator (M. Pepper, University of Washington). 4get-GFP and ICAM-1.KO mice were crossed to OT-II to generate 4get-GFP OT-II and ICAM-1.KO OT-II lines respectively.

Six-to twelve-week-old male and female mice were kept in specific pathogen-free (SPF) conditions at an Association for Assessment and Accreditation of Laboratory Animal Care-accredited facility at the University of Washington, South Lake Union campus. All procedures were approved by the University of Washington Institutional Animal Care and Use Committee.

### Adoptive transfers

For adoptive transfers, naïve CD45.1^+^ OT-II, CD45.1^+^CD45.2^+^ OT-II, or CD45.2^+^ 4get-GFP OT-II T cells were isolated from LNs and spleens using the naïve CD4^+^ T cell isolation kit (Miltenyi Biotec). The average purity of OT-II cells was approximately 75-80% for all experiments. 1×10^6^ (unless otherwise noted) naïve OT-II cells were transferred into hosts intravenously via retro-orbital injection 1-3 days prior to immunization.

### Immunizations and blocking antibodies

The following adjuvants and amounts per immunization site were used: 20µg of CpG ODN 1668 (AdipoGen), Alhydrogel (“Alum”) (Invivogen) diluted 1:2 with PBS, Dibutyl pthalate (“DBP”) (Sigma Aldrich) diluted 1:1 with 100% Acetone, 50µg Papain (Sigma-Aldrich), and 10µg Endotoxin-free Ovalbumin (“OVA”) (Invivogen). In some studies, the dose of OVA given ranged from 0.1-100µg per injection. For antigen presentation studies, 10μg of LPS-free EαGFP (gift from M. K. Jenkins, University of Minnesota) plus 50μg of papain were injected intradermally in the ear pinnae or the forepaw. Adjuvants were mixed with PBS in a 20µl total volume per immunization site and injected in the hind or front footpads, subcutaneously in the flank skin, at the tail base, or intradermally in the ear pinnae (to target the popliteal, Brachial, Inguinal, and Auricular dLNs, respectively) as indicated. For some studies, DBP was “painted” onto ear or footpad skin by pipetting 20µl volume slowly, allowing drying between drops with the mouse under anesthesia until liquid was completely dried.

For *in vivo* antibody blocking studies, 100µg of anti-CD62L (clone Mel-14, BioXcell) was injected intraperitoneally 6 hours post immunization. In some studies 100µg anti-LFA-1α (clone M17/4, BioXcell), 100µg rat IgG2a isotype control (BioXcell), 100µg anti-CD28 (clone 37.51, BioXcell), 100µg polyclonal Syrian hamster IgG (BioXcell), or 100µg anti-IL-2 (clone JES6-1A12, BioXcell) was injected intraperitoneally 24 hours post immunization.

### Confocal microscopy

For confocal imaging, PFA-fixed and sectioned LN tissues were imaged as previously described using a Leica SP8 microscope^58^. Briefly, isolated LN tissues were fixed using BD Cytofix (BD Biosciences) diluted 1:3 in PBS for 24 hours at 4°C then dehydrated with 30% sucrose solution for 24-48 hours at 4°C. LNs were then embedded in OCT compound (Tissue-Tek) and stored at -20°C. LNs were section on a Thermo Scientific Micron HM550 cryostat into 20µm sections and stained as previously described^58^. A Leica SP8 tiling confocal microscope equipped with a 40x 1.3NA oil objective was used for image acquisition. All raw imaging data was processed and analyzed using Imaris (Bitplane).

### Histo-cytometry and CytoMAP

Histo-cytometry analysis was performed as previously described^38, 58^. Briefly, multiparameter confocal images were corrected for fluorophore spillover using the built-in Leica Channel Dye Separation module. Single stained controls were acquired using UltraComp eBeads (Invitrogen) that were incubated with fluorescently conjugated antibodies, mounted on slides with Fluormount-G slide mounting media (ThermoFisher), and imaged. All images were visualized and analyzed using Imaris (Bitplane). For analysis of myeloid cells, a combinatorial myeloid channel was created by adding normalized signals for CD11c, MHC-II, CD207, CD301b, and Sirpα using the Imaris XT channel arithmetic module, and this sum myeloid channel was used for myeloid isosurface object creation. For T cells, a combined activated T cell channel was created by adding normalized signals for Ki67 and IRF4; T cell isosurface objects were next created on this channel and then further gated on the congenic CD45.1 or CD45.2 signal. In all analyses, object statistics were exported to FlowJo software (FlowJo, LLC) for gating and phenotypic characterization. T-B border regions were manually created using B220 or MHC-II (B cell follicles) and CD3 (T cell zone) staining and represented as a surface. T-B border localization was calculated as the frequency of OT-II cells within the T-B border surface region. Clustering analysis was performed by creating isosurfaces on CD45.1 congenic signal without cell splitting. Surfaces were then filtered by volume and surfaces exceeding 1400µm^3^ were added and presented as a ratio of macro-clustered surface volume (greater than 1400 µm^3^) to total OT-II surface volume. Spatial correlation analysis was performed in CytoMAP^42^. In brief, the position of all myeloid and T cell objects within LNs was used for virtual raster scanning with 50-µm radius neighborhoods. The Pearson correlation coefficient was calculated for the number of cells of the different cell types within these neighborhoods. Raster-scanned neighborhoods were also used for clustering based on cell type abundance to identify distinct region types, and these regions were used for heatmap and positional visualization of regions in dLNs.

### Cell isolation and flow cytometry

For myeloid cell analysis, LN tissues were mechanically disrupted and subject to digestion in PBS with 10% fetal bovine serum (FBS) with DNase I (100µg/mL; Sigma), Dispase II (800µg/mL; Sigma), and Collagenase P (200µg/mL; Sigma) at 37°C shaking at 150rpm for 30 minutes with periodic manual disruption. Flow cytometric studies of T cells in lymph nodes did not use enzymatic digestion. In some studies, mice were injected intravenously with 1µg of anti-Thy1.2-BUV395 (clone 30-H12; BD Biosciences) ∼5 minutes prior to sacrifice. Lung tissue was digested in complete RPMI with Liberase (70µg/mL; Roche) and Aminoguanidine (10mM; Sigma) and tissue was dissociated on the gentleMACS Dissociator (Miltenyi Biotec) as previously described^22^. Cell staining was conducted in the presence of Fc Block (2.4G2, Tonbo Biosciences) at 4°C for 30 minutes for all surface markers except CXCR5-biotin which was stained at room temperature for 45 minutes. Intracellular staining was performed for 45 minutes at 4°C after fixation with the FOXP3 Fix/perm kit (Invitrogen). In some studies, an addition permeabilization step was performed in 90% ice-cold methanol prior to intracellular staining. Data were acquired on an Aurora flow cytometer (Cytek) and analyzed using FlowJo software (BD Biosciences).

### CD4^+^ T cell isolation and culture

For *in vitro* T cell stimulation experiments, naïve CD4^+^ T cells were isolated from LNs and spleens using the naïve CD4^+^ T cell isolation kit (Miltenyi Biotec) into complete RPMI media. 400,000 cells were then plated on pre-treated plates coated with the indicated concentrations of anti-CD3ԑ (145-2C11, Thermo Scientific) and anti-CD28 (37.51, Thermo Scientific) under Th0, +anti-IFNγ 10ug/mL; (XMG1.2, Biolegend), or Th2 (anti-IFNγ 10µg/mL (XMG1.2, Biolegend), rIL-4 50ng/mL (Peprotech)) conditions. Cells were cultured for 48 hours at 37°C and 5% CO_2_ before fixation and flow cytometric analysis.

### RNA sequencing

Single cell suspensions from tissues were prepared as described above for myeloid cells. Cells were sorted from pooled dLNs from 3 individual mice for each group. 500 of each cDC2 cell type was sorted on an Aria III (BD Biosciences) directly into reaction buffer from the SMART-Seq v4 Ultra Low Input RNA Kit for Sequencing (Takara), and reverse transcription was performed followed by PCR amplification to generate full length amplified cDNA. Sequencing libraries were constructed using the NexteraXT DNA sample preparation kit with unique dual indexes (Illumina) to generate Illumina-compatible barcoded libraries. Libraries were pooled and quantified using a Qubit Fluorometer (Life Technologies). Sequencing of pooled libraries was carried out on a NextSeq 2000 sequencer (Illumina) with paired-end 59-base reads, using NextSeq P2 sequencing kits (Illumina) with a target depth of 5 million reads per sample. Base calls were processed to FASTQs on BaseSpace (Illumina), and a base call quality-trimming step was applied to remove low-confidence base calls from the ends of reads. The FASTQs were aligned to the GRCm38 mouse reference genome, using STAR v.2.4.2a and gene counts were generated using htseq-count. QC and metrics analysis was performed using the Picard family of tools (v1.134). Further downstream analysis was performed using publicly available RNAseq toolkits. The Degust toolkit^104^ (v4.1.1) with integrated Voom/Limma R package was used for differentially expressed gene analysis and generation of volcano plots. Only genes with count per million (CPM) ≥ 10 were analyzed further. Genes were filtered based on a false discovery rate cutoff ≤ 0.05 and a minimum expression fold change ≥ 2. DEGs were input into the WebGestalt gene set analysis toolkit^105^ to identify Biological Processes Gene Ontology (GO) terms and generate the associated graphs. Heatmap data tables were generated by inputting DEG data into the BIOMEX toolkit^106^. PCA plots were generated via the DEBrowser toolkit^107^ (v1.26.3), where only genes with CPM ≥ 10 were analyzed.

### Antibodies & staining reagents

Antibodies used for staining sections for confocal imaging or isolated cells for flow cytometry include: CD64 (clone X54-57.1; BioLegend), B220 (clone RA3-6B2; Biolegend), SIRPα (clone P84; BD Biosciences), CD11c (clone N418; BD Biosciences), CD11b (clone M1/70; Biolegend), MHCII (clone M5/114.15.2; BioLegend), IRF4 (clone IRF4.3E4; BioLegend), CD45.1 (clone A20; BioLegend), Ki67 (clone B56; BD Biosciences), Y-Ae (clone eBioY-Ae; ThermoFisher Scientific), Thy1.2 (clone 30-H12; BD Biosciences), CD3 (clone 17A2; BD Biosciences), Tbet (clone 4B10; BioLegend), NK1.1 (clone PK136; BioLegend), CD19 (clone 6D5; BioLegend), CD44 (clone IM7; BioLegend), XCR1 (clone ZET; BioLegend), GATA3 (clone L50-823; BD Biosciences), CXCR3 (clone CXCR3-173; BioLegend), CXCR5 (clone 2G8; BD Biosciences), pS6 (clone 2F9; Cell Signaling Technologies), PD-1 (clone RMP1-30; BioLegend), CD45.2 (clone 104; BioLegend), CD25 (clone PC61; ThermoFisher Scientific), EpCAM (clone G8.8; ThermoFisher Scientific), CD103 (clone M290; BD Biosciences), CD301b (clone URA-1; BioLegend), CD80 (clone 16-10A1; ThermoFisher Scientific), CD86 (clone GL1; BD Biosciences), PDL-1 (clone MIH5; ThermoFisher Scientific), PDL-2 (clone MIH37; BD Biosciences), BCL6 (clone K112-91; BD Biosciences), BATF (clone D7C5 rabbit; Cell Signaling Technologies), Anti-GFP (goat; Novus Biologics), ICAM-1 (clone 3E2; BD Biosciences), CD207 (clone 929F3.01; Dendritics), CD4 (clone GK1.5; BD Biosciences), huCD2 (clone RPA-2.10; ThermoFisher Scientific), pSTAT5 (clone 47/STAT5(pY694); BD Biosciences), pSTAT6 (clone J71-773.58.11; BD Biosciences), LIVE/DEAD Near-IR (ThermoFisher), Chicken anti-goat (ThermoFisher) and Donkey anti-rabbit (ThermoFisher).

### Statistics

Statistical analysis was performed using GraphPad Prism software. The statistical significance of differences in mean values between two groups was analyzed by a two-tailed unpaired student’s *t* test with Welch’s correction. Paired *t* tests were performed when comparing responses within the same experimental tissue. In bar graphs for all figures, data is shown as mean with standard deviation. ****p<0.0001; ***p<0.001; **p<0.01; *p<0.05; p>0.05 not significant (ns). Unless otherwise noted, all data points represent independent LNs.

## Acknowledgments

We thank current and former lab members A. de la Cruz, E. Cheng, J. Huang, J. Leal, and J. Chao, for help with conceptualizing and conducting experiments. We also thank J. von Moltke and J. McGinty for help with *Nippostrongylus brasiliensis* infections. Additionally, we thank M. Pepper, S. Ziegler, E. Tait Wojno, and J. von Moltke for additional mice, reagents, and resources.

## Author contributions

M.R.L-C and M.Y.G conceptualized the study. M.R.L-C performed all experiments and analyzed data. E.A.S analyzed RNA sequencing data. M.R.L-C and M.Y.G wrote, edited, and reviewed the manuscript. M.Y.G. supervised the project.

